# Cell-type specific asynchronous modulation of PKA by dopamine during reward based learning

**DOI:** 10.1101/839035

**Authors:** Suk Joon Lee, Bart Lodder, Yao Chen, Tommaso Patriarchi, Lin Tian, Bernardo L. Sabatini

## Abstract

Canonical reinforcement learning models postulate that dopamine neurons encode reward prediction error (RPE) and provide a teaching signal to striatal spiny projection neurons (SPNs) in the form of dopamine (DA) release. DA is thought to guide learning via dynamic modulation of protein kinase A (PKA) in SPNs. However, this fundamental assumption remains untested in behaving animals. Here we utilized multi-channel fiber photometry and fluorescence lifetime photometry (FLiP) to monitor the activity of DA neurons, extracellular DA levels, and net PKA activity in SPNs in the nucleus accumbens during learning. We found dynamic encoding of RPE in the activity of DA neurons, which is both necessary and sufficient to explain striatal DA levels and SPN PKA activity. The modulation of PKA in SPNs that express type-1 (D1R-SPNs) and type-2 (D2R-SPNs) DA receptors was dichotomous such that in each cell class it is selectively sensitive to increases and decreases in DA, respectively, and occur at and support different phases of learning. Thus, PKA-dependent pathways in D1R- and D2R-SPNs are asynchronously engaged by RPE-encoding DA signals to promote different aspects of reinforcement learning: the former responsible for the initial association between action and outcome and the latter responsible for refining the learned association.

## Introduction

Across phylogeny, dopamine (DA) release in the brain induces cellular plasticity pathways that underlie learning and behavioral adaptation^1^. In mammals, DA action in the nucleus accumbens (NAc), a ventral part of striatum that is heavily innervated by ventral tegmental area (VTA) DA neurons, is thought to mediate the association of motor actions with good or bad outcomes such that an individual repeats or avoid behaviors that previously lead to good or bad outcomes, respectively^2,3,4,5^. Supporting this model, manipulations of the activity of VTA DA neurons and NAc DA levels established the sufficiency of DA release in the NAc in reinforcing an action^6,7,8,9^. In fact, the reinforcing effect of DA in the NAc is thought to be the mechanism of addiction for many drugs of abuse^10^. Additionally, involvement of DA in reinforcement learning is further supported by many studies^11,12,13,14,15,16,17^ showing that the activity of VTA DA neurons and NAc DA levels encode reward prediction error (RPE) - the difference between the value of the actual and expected outcome of an action. In computational models, RPE is a useful teaching signal for forming an association between a past action and a present outcome^11,18^.

What is the mechanism of DA action in the NAc? The anatomical and molecular organizations of spiny projection neurons (SPNs), the principle cells of the NAc, suggest an antagonistic model of DA action on SPNs during reward based learning^3,19^. NAc SPNs, analogous to the direct and indirect pathway SPNs of the dorsal striatum, are divided into striatomesencephalic SPNs, which innervate various midbrain regions, and striatopallidal SPNs, which innervate the ventral pallidum^20,21^. This anatomic division correlates with transcriptional and biochemical differences: striatomesencephalic SPNs express G_αs_-coupled type 1 DA receptors (D1Rs) by which DA enhances cAMP production and protein kinase A (PKA) activity whereas striatopallidal SPNs express G_αi_-coupled type 2 DA receptors (D2Rs) by which DA inhibits cAMP production and suppresses PKA activity^20^. Models of reinforcement learning^3^ incorporate these differences to link RPE-reflecting DA signals to the differential modulation of excitability^22,23^, synaptic plasticity^24^, and transcription via PKA in each cell type^25,26^.

Despite the extensive work on DA and the NAc described above, how complex patterns of DA neuron activity translate into the dynamic modulation of PKA in SPNs *in vivo* is still unknown due to difficulty of monitoring intracellular signaling in behaving animals. D1R- and D2R-expressing SPNs also express many other classes of GPCRs^27^ as well as Ca^2+^-dependent adenylyl cyclases^28^ that may also regulate PKA during behavior and potentially dominate DA-specific effects. Also, the transformation between the dynamics of NAc DA and intracellular PKA activity could be shaped by the differential affinities of D1Rs and D2Rs^29,30^ expressed by each SPN class. Further complicating the relationship between DA neuron activity and SPN PKA activity, a recent study indicated that the soma activity of DA neurons do not necessarily correlate with DA levels in the region they innervate^17^.

Here, we investigate these unknowns by measuring VTA DA neuron Ca^2+^ levels, NAc DA levels, and net PKA activity in SPNs in freely behaving mice while they learn a locomotor task performed for food rewards. Simultaneous measurements of VTA DA neuron bulk Ca^2+^ levels (referred as DA neuron activity below for simplicity) and NAc DA levels by multi-channel fiber photometry indicate that they are correlated with each other and consistent with RPE. Using fluorescence lifetime photometry (FLiP), which permits dynamic measurement of intracellular biochemical signals in genetically-defined neurons using a PKA substrate sensor (FLIM-AKAR) ^31^, we monitored endogenous net PKA activity (the balance between PKA and phosphatase activity) in D1R- and D2R-expressing SPNs (D1R-SPNs and D2R-SPNs, respectively)^32^. We reveal dynamic changes in D1R-SPN and D2R-SPN PKA activity that are asynchronously and differentially triggered during reinforcement learning. These changes in PKA activity are explained by the patterns of VTA DA neuron activity and NAc DA levels. Furthermore, combining FLiP with pharmacological and optogenetic manipulations, we demonstrate both the necessity and sufficiency of transient changes in DA in modulating SPN PKA activity. Lastly, we reveal that PKA activity in D1R- and D2R-SPNs are responsible for different phases of learning by investigating the effects of selective inhibition of PKA in D1R- and D2R-SPNs. In summary, our results support a revised dichotomous model of DA and basal ganglia dependent learning mechanisms in which PKA in D1R- and D2R-SPNs is asynchronously engaged to mediate the action of RPE-encoding DA signals during learning.

## Results

### Fluorescence lifetime photometry (FLiP) reveals bidirectional changes in SPN PKA activity *in vivo*

To monitor SPN PKA activity in vivo, we performed fluorescence lifetime photometry of FLIM-AKAR through an optical fiber implanted in the NAc^32^. Briefly, FLIM-AKAR is a Förster resonance energy transfer (FRET)-based PKA activity reporter consisting of a donor fluorophore and a dark acceptor^31^. Phosphorylation of the sensor by PKA induces a conformational change that increases FRET efficiency between the donor and the acceptor, resulting in faster fluorescence decay of the donor. Therefore, as previously validated *in vitro*^31,33^ and *in vivo*^32^, a reduction (negative change) in fluorescence lifetime of FLIM-AKAR represents an increase in the net phosphorylation of PKA substrates in cells expressing the sensor.

To examine the ability of each DA receptor class to dynamically modulate PKA in awake animals, we used FLiP to monitor net PKA activity in SPNs following intraperitoneal (IP) delivery of DA receptor agonists and antagonists. We expressed FLIM-AKAR in either D1R- or D2R-SPNs in the NAc by injecting an adeno-associated virus (AAV) that expressed the sensor in a Cre-dependent manner (AAV1-FLEX-FLIM-AKAR) into *Drd1a-Cre* or *Adora2a-Cre* mice, respectively (Fig. 1a). Fluorescence was detected using an optical fiber that was implanted above the injection site and the optical system described in the Extended Data Fig. 1. Fluorescence lifetimes were measured with time correlated single photon counting, and a lifetime histogram (Fig. 1a, Extended Data Fig. 2a) was produced every second resulting in a 1 Hz measurement of average fluorescence lifetime.

**Fig. 1.**
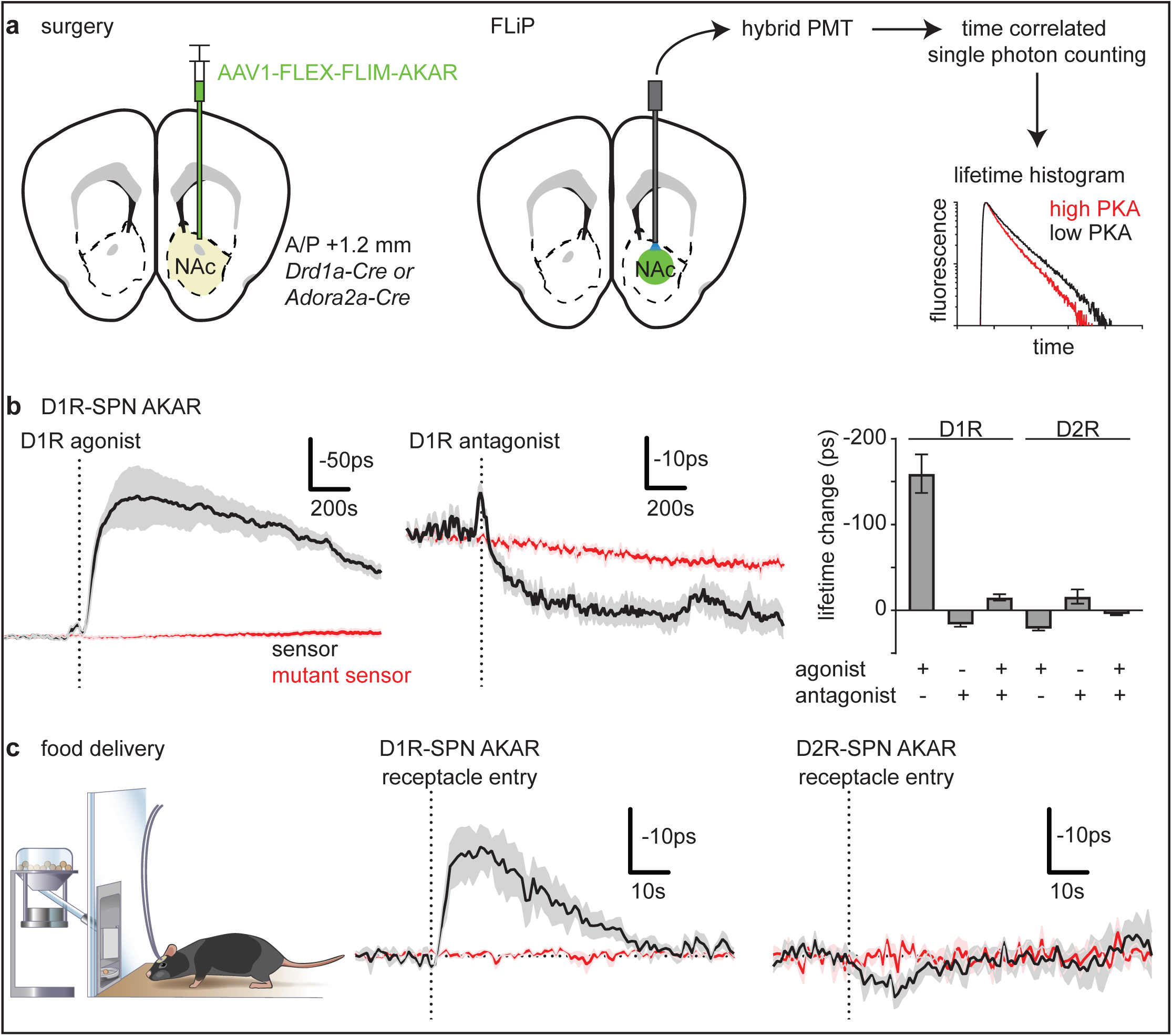
Fluorescence lifetime photometry (FLiP) reveals bidirectional changes in SPN PKA activity *in-vivo*. **a)** Schematic of a coronal section at 1.2 mm depicting viral injection of AAV1-FLEX-FLIM-AKAR into the NAc of *Drd1a-Cre* or *Adora2a-Cre* mice (*left*). An optical fiber was implanted 200 μm above the injection site (*middle*). A hybrid PMT and time correlated single photon counting were used to measure the fluorescence lifetime of FLIM-AKAR, which provides an estimate of the ratio between phosphorylated and un-phosphorylated FLIM-AKAR (*right*). Higher PKA activity results in a faster fluorescence decay (lower fluorescence lifetime) of the sensor (red). **b)** Average changes in fluorescence lifetime of FLIM-AKAR expressed in D1R-SPNs of *Drd1a-Cre* mice that were injected with D1R agonist (*left*) or antagonist (*right*) (n=7 mice each). The dashed vertical lines indicate the times of the end of the injection. For FLIM-AKAR (black), a decrease in lifetime represents an increase in phosphorylation by PKA. For clarity, all fluorescence lifetimes are plotted inverted such that upward transients indicate increased PKA phosphorylation. A similar measurement was made using FLIM-AKAR^T391A^ (red), which has a point mutation at the PKA phosphorylation site (n=7 mice). In this and all average responses, the solid line and shaded region show the average +/− SEM across mice. **c)** Summary of peak lifetime changes of FLIM-AKAR in D1R-SPNs of *Drd1a-Cre* mice following D1R agonist and/or antagonist injections (n=7 mice each) or those in D2R-SPNs of *Adora2a-Cre* mice following injection of D2R agonists and/or antagonists (n=6 each except n=5 for D2R antagonist + agonist trials). Error bar=SEM of averages across mice. **d)** *left*, Schematic of a mouse receiving food from an automated dispenser while FLIP measurements are made through a fiber optic. Fluorescence lifetime changes of FLIM-AKAR expressed in D1R-SPNs of a *Drd1-Cre* mouse (*middle*) or D2R-SPNs of an *Adora2a-Cre* mouse (*right*) that was given a food reward (dashed vertical line = time point of food pellet receptacle entry). Mice were food restricted at least one day before measurement of the food reward response. Black=FLIM-AKAR. Red= FLIM-AKAR^T391A^. For each mouse, data were collected across 10 trials. n=4 mice (D1R-SPN FLIM-AKAR, sensor), 3 mice (D1R-SPN FLIM-AKAR, mutant sensor), 4 mice (D2R-SPN FLIM-AKAR, sensor), 4 mice (D2R-SPN FLIM-AKAR, mutant sensor).

In *Drd1a-Cre* mice, FLIM-AKAR expressed in D1R-SPNs in the NAc reported an increase in net PKA activity (~160ps lifetime change) following systemic administration of the D1R agonist SKF 81297 hydrobromide (10 mg/kg IP) (Fig. 1b, Extended Data Fig. 2a), consistent with D1R-mediated activation of PKA in these cells. The D1R-agonist induced change was not observed for D1R-SPNs expressing mutant AKAR (FLIM-AKAR^T391A^), which has a point mutation at its PKA phosphorylation site. In contrast, the D1R antagonist SKF 83566 hydrobromide (3mg/kg IP) resulted in a small (~17ps) but significant increase FLIM-AKAR lifetime (Fig. 1b), indicating a slight reduction in D1R-SPNs’ net PKA activity. Furthermore, pre-administration of the D1R antagonist largely blocked the D1R agonist response (Fig. 1b), confirming the specificity of the agonist action on D1Rs.

To assay the modulation of PKA by DA receptors (D2R) in D2R-expressing SPNs, we used *Adora2a-Cre* mice and the Cre-dependent viruses as above. D2R agonist sumanirole maleate (4mg/kg IP) significantly increased (~22ps) the fluorescence lifetime, indicating suppression of D2R-SPNs’ net PKA activity (Fig. 1b, Extended Data Fig. 2b). Conversely, the injection of D2R antagonist (eticlopride hydrochloride, 0.5mg/kg IP) decreased FLIM-AKAR fluorescence lifetime (~16ps), demonstrating an increase in D2R-SPNs’ net PKA activity (Fig. 1b, Extended Data Fig. 2b). Furthermore, pre-injection of D2R antagonist blocked the effect of the D2R agonist (Fig. 1b), and no effects were seen in phosphorylation-site mutant AKAR expressing mice. In addition, we found that neither IP nor intraventricular (IV) injection of sulpiride, a reported D2R antagonist, significantly changes D2R-SPNs’ net PKA activity, whereas IV but not IP injection of an A2A adenosine receptor agonist (CGS 21680 hydrochloride) increases D2R-SPNs’ net PKA activity (Extended Data Fig. 2d and e).

Thus, pharmacological modulation of DA receptors in D1R- and D2R-SPNs results in large and prolonged changes in net PKA activity that are detectable by FLiP. In addition, antagonism of DA receptors revealed a small basal tone of PKA activity in both cell types, suggesting that behaviorally-induced up and down regulation of DA has the potential to dynamically regulate PKA in both cell classes. Indeed, the ability to detect behaviorally induced changes in SPNs PKA activity is suggested in the above experiments: a small (~10ps) transient (~70s) reduction in FLIM-AKAR fluorescence lifetime was observed in D1R-SPNs at the time point of IP injection independent of the nature of drug administered (agonist, antagonist, or control saline) (Fig. 1b; Extended Data Fig. 2b). Since it persisted in the presence of D1R-antagonists but was not present in mutant AKAR expressing mice (Extended Data Fig. 2b), this minor change likely reflects activation of D1R-SPN PKA following scruffing and injection that is potentially independent of D1R activation. Furthermore, as this change was not seen in D2R-SPNs (Extended Data Fig. 2c), this transient indicates the potential for differential PKA signaling in each class of SPN induced by environmental stimuli.

In order to further observe potential modulation of SPN PKA activity in response to natural stimuli, we examined fluorescence lifetime responses to unexpected food delivery in food-restricted *Drd1a-Cre* and *Adora2a-Cre* mice expressing FLIM-AKAR in D1R- and D2R-SPNs, respectively (Fig. 1c). Delivery of food pellets in a receptacle significantly reduced fluorescence lifetime (~20ps) of FLIM-AKAR in D1R-SPNs. The food induced response was much shorter-lived (~40-60s) than the response due to D1R agonists (Fig. 1c), potentially reflecting PKA activation induced by a brief increase in DA at the time of reward. Conversely, food reward induced a small but statistically significant increase (~5ps) in fluorescence lifetime in D2R-SPNs mice, demonstrating a weak suppression of net PKA activity in these cells at the time of reward (Fig. 1c). Both responses were absent in PKA-phosphorylation site mutant FLIM-AKAR (FLIM-AKAR^T391A^) expressing neurons.

In summary, SPN PKA activity manipulations by pharmacological manipulation of DA receptors and natural stimuli demonstrate the capacity of FLiP to detect bidirectional changes in PKA activity *in vivo* as well as the DA receptor dependent regulation of PKA activity in SPNs. In addition, the pharmacology experiments with DA receptor antagonists provide an *in vivo* evidence for the effect of basal DA levels on D1Rs and D2Rs. However, possible inverse agonist actions of the antagonists cannot be ruled out.

### Plasticity of DA release and DA neuron activity dynamics across learning is consistent with RPE

Since fluorescence lifetime changes of FLIM-AKAR in response to a food reward can last up to 60s, we designed a slow time-scale operant conditioning task (Fig. 2a) to investigate how DA neuron activity, NAc DA, and SPN PKA activity are modulated throughout the process of learning. Mice were habituated (i.e. given 10 free pellets) to the arena for 1 day (day 0) and then trained on the full task for 11 days (day 1-11). Most mice reached 90% success rate in 9 days (Extended Data Fig 3a.). Mice increased their success rate by reducing the latency to enter the receptacle zone as well as by staying longer in the receptacle zone (Extended Data Fig. 3a). Furthermore, they learn to stay in the trigger zone in order to initiate trials quickly after the unindicated minimum 120s inter-trial interval (Extended Data Fig. 3a). On day 12, the reward was omitted from 25% of successful trials to collect “reward omission” trials. On day 13, the LED was turned off for the entire session in order to collect “LED omission” trials in which mice occasionally managed to perform the correct movements despite the absence of the LED cue and thus received “unexpected” rewards.

**Fig. 2.**
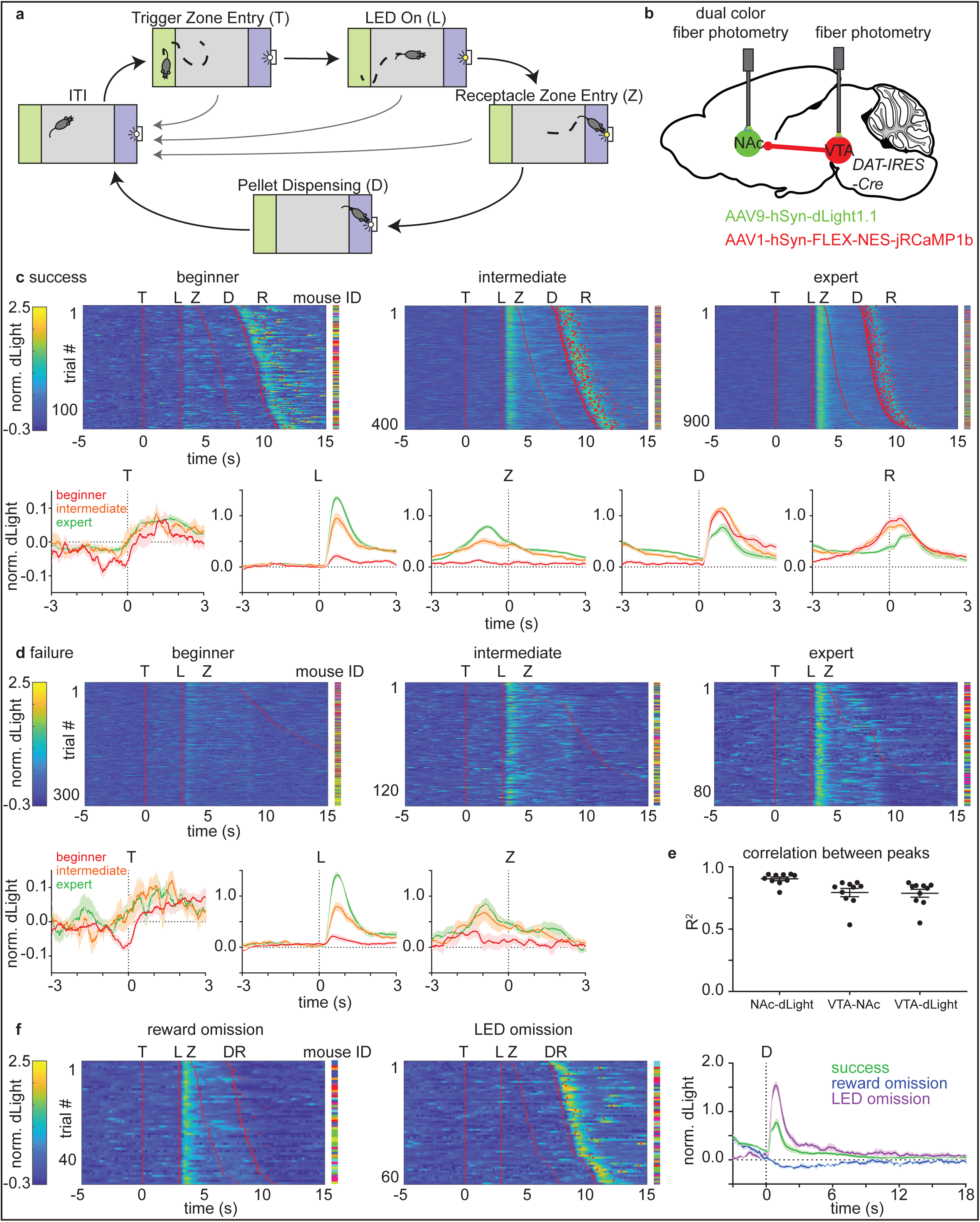
Plasticity of DA release and DA neuron activity dynamics across learning is consistent with RPE. a) Schematic of the behavioral paradigm. After the inter-trial interval (ITI, minimum 120s), a mouse can initiate a trial by entering (T) and staying in the trigger zone (green) for 3 consecutive seconds. Successful initiation of a trial is signaled by activation of an LED (L). After the LED turns on, the mouse must enter the receptacle zone (blue) within 5s (Z) and stay in this zone for 3 consecutive seconds to trigger dispensing of a food reward (D). Failure to successfully accomplish each step returns the apparatus to the ITI, and the mouse must initiate a new trial. The LED turns off at the end of each trial whether by success (food delivery) or failure (return to the ITI). Each daily session consists of 20 self-initiated trials. b) Schematic of a sagittal section depicting viral expression and fiber implantation in *DAT-IRES-Cre* mice. AAV9-hSyn-dLight and AAV1-hSyn-FLEX-NES-jRCaMP1b were injected into NAc and VTA, respectively. Fibers were implanted 200μm above the two injections sites. The NAc fiber was used to collect both dLight and DA neuron terminal jRCaMP signals. The VTA fiber was used to collect DA neuron somatic jRCaMP signal. c) dLight response in success trials during learning. Top: heatmaps of dLight responses (df/f normalized across all sessions for each mouse such that the 99 percentile response = 1). Each row represents an individual trial. Red lines or dots indicate behavioral time stamps (T=trigger zone entry, L=LED on, Z=receptacle zone entry, D=pellet dispensing, R=receptacle entry). The mouse ID for each row is represented by a different color. n=117 trials (beginner), 400 trials (intermediate), 965 trials (expert) from 10 mice. Bottom: average dLight responses (normalized df/f) across mice aligned to each behavioral time stamps. The average responses for beginner, intermediate, and expert trials are shown in red, orange, and green, respectively. Shaded area=SEM of averages across mice. d) As in panel (c) for dLight responses in failure trials (failure to enter the receptacle zone and failure by prematurely exiting the receptacle zone) during learning showing heatmaps (top, n=339 beginner, 136 intermediate, and 89 expert trials from 10 mice) and average dLight responses (bottom) across mice aligned to each behavioral time stamp. e) Correlations between peak amplitudes of signals that rise at least 2 SD above baseline (measured in the −20s before trigger zone entry) across all signal pairs (NAc-dLight: DA neuron terminal jRCaMP and dLight signal; VTA-NAc: DA neuron soma and terminal jRCaMP signal; VTA-dLight: DA neuron soma jRCaMP and dLight signal). Each dot is the R^2^ value for data from a single mouse average across sessions. Error bar = SEM of averages across mice. n=10 mice. f) dLight responses as above in reward omission trials (*left*) and rewarded LED omission trials (*middle*) for expert mice. n=50 trials (reward omission), 62 trials (LED omission) from 10 mice. *right*, Average dLight response of expert (trained) mice aligned to the time of pellet dispensing for success trials (green), reward omission trials (blue), rewarded LED omission trials (purple).

Given the recent finding that NAc core DA release can be dissociated from increases in VTA DA neuron spiking^17^, we simultaneously monitored somatic DA neuron activity in the VTA as well as the activity of VTA DA neuron axons and ensuing DA transients in the NAc. We tested if the three signals encode reward prediction error (RPE) in freely moving mice performing the behavior task. To simultaneously monitor both DA neuron activity and NAc DA release, we expressed jRCaMP1b^34^ in VTA DA neurons in a Cre-dependent manner by injecting AAV1-hSyn-FLEX-NES-jRCaMP1b into *DAT-IRES-Cre* mice and expressed dLight^35^ in the NAc neurons by injecting Cre-independent AAV9-hSyn-dLight1.1 (Fig. 2b) into NAc. We implanted optical fibers into both VTA and NAc and used the former to monitor VTA DA neuron somatic activity and the latter to monitor both VTA DA neuron terminal activity and DA release in the NAc. A highly sensitive light detector and optimized light collection allowed us to use a relatively low level of excitation light (5~16 µW) and thereby perform fiber photometry in all sessions across 13 consecutive days to track progressive changes in neural signals (Extended Data Fig. 1).

In beginner animals, NAc DA levels increased rapidly following reward delivery whereas only minimal changes in DA at the time of activation of the LED cue were observed (Fig. 2c-d). In expert animals, the magnitude of DA release after reward was lower than that of beginner animals, whereas there was a large release of DA triggered by the LED cue (Fig. 2c-d). Furthermore, as evidenced by intermediate performance states (i.e. intermediate animals), the shift in DA release from the reward to the cue occurred gradually across training, a pattern seen in both individual mice and averages across mice (Fig. 2c-d, Extend Data Fig. 3c-d), which is consistent with the classic RPE theory. In addition, the significant reduction of reward-associated responses across training was not explained by photo-bleaching of dLight (Extended Data Fig. 3d). Lastly, DA release patterns in unrewarded failure trials, including both failures to enter the receptacle zone on time and failures by prematurely exiting the receptacle zone after entering it, showed LED response similar to that in success trials but lacked reward responses at the time point of receptacle entry (Fig. 2d).

We further tested if the pattern of DA release in this learning paradigm is consistent with RPE by examining reward omission (day 12) and LED omission (day 13) trials. Consistent with previous studies, DA levels dipped below baseline at the time of expected reward delivery when the reward was omitted (Fig. 2f). In LED omission trials, success rate plummeted, suggesting that even trained animals rely on the LED cue to perform the task and are not simply performing a timing task (Extended Data Fig. 3a). Furthermore, there was no significant increase in DA level at the time of an omitted LED cue, indicating that the DA response when the LED turns on was truly driven by this external cue rather than an internal expectation (Extend Data Fig 3b). In addition, the DA peak after reward was larger in rewarded LED omission trials than in regular trials (Fig. 2f), confirming that reward expectation (or lack of) bi-directionally modulates DA levels.

The patterns of DA during the behavioral task also existed in DA neuron soma and terminal activity patterns (Extended Data Fig. 3b-d). We formally tested the relationship across the signals by calculating correlation between the sizes of peaks in each signal that rise 2 standard deviations above baseline (Fig. 2e), and found a tight correlation (R^2^ >= ~0.8). To compare signals with different kinetics across all time points and avoid spurious correlations introduced by slow fluorescent reporters, we assumed that each signal follows an autoregressive process (see Methods) and deconvolved the fluorescence transients to extract population events. Even for the comparison across all time points, DA release events (as well as DA terminal activity) and DA soma activity remained well correlated (R^2^ ~0.5) (Extended Data Fig. 4b). Furthermore, we created a generalized linear model that relates the behavioral parameters to each fluorescence signal (Extend Data Fig. 5a) and found that, across all stages of learning, the models were similar for all three fluorescence signals (Extended Data Fig. 5b-c). Overall, these results indicate that NAc DA levels during the behavioral task are strongly modulated by DA neuron soma activity, which carries a characteristic of RPE signal. In addition to these rapid patterns of DA release, small long-lasting changes in baseline and peak DA levels were seen across trials indicating additional complex effects of trial history on NAc DA (Extended Data Fig. 6).

To control for potential movement and hemodynamic artifacts, we performed similar photometric analysis in mice expressing eGFP or mutant dLight that has a mutation in the DA binding site (Extended Data Fig. 7a). Interestingly, after learning there was a small (more than 10 fold smaller than the actual sensor signals) transient reduction in the NAc fluorescence of both eGFP and mutant dLight that lasts ~30s, suggesting a possible hemodynamic effect at the time of the learned cue (Extended Data Fig. 7b). Furthermore, analysis of a separate cohort of mice expressing only a single sensor (Extended Data Fig. 7c) confirmed the lack of significant optical crosstalk during the simultaneous green and red photometry.

### D1R- and D2R-SPN PKA activity are dynamically modulated by DA at different stages of learning

DA release in the two brain hemispheres are symmetric in our non-lateralized behavior task (Extended Data Fig. 8a-b). Therefore, we performed simultaneous measurements of DA levels and net PKA activity in SPNs by expressing dLight and FLIM-AKAR in different hemispheres (Fig. 3a). Consistent with dLight / jRCaMP experiments discussed above, the dLight response aligned to the cue increased whereas that aligned to the reward decreased across training (Fig. 3b). Following the patterns of DA release, in beginner animals, D1R-SPNs’ net PKA activity increased at the time of reward consumption whereas, in expert animals, it began to increase at the time of the LED cue (Fig. 3b and Extended Data Fig. 9). For rewarded trials in trained animals, the change of net PKA activity detected by the sensor likely integrated the effect of both cue- and the reward-evoked DA release. Nevertheless, it was still possible to separate out the contribution of the cue- and the reward-evoked PKA activity by comparing success and failure trials, which demonstrated that D1R-SPN PKA is activated by both the LED cue and the reward in expert animals (Fig. 3b-c). Furthermore, D1R-SPN PKA activation was larger in the rewarded LED omission trials than in the regular rewarded trials for expert animals, consistent with the larger DA release induced by unexpected reward triggering greater PKA activation (Fig. 3c).

**Fig. 3.**
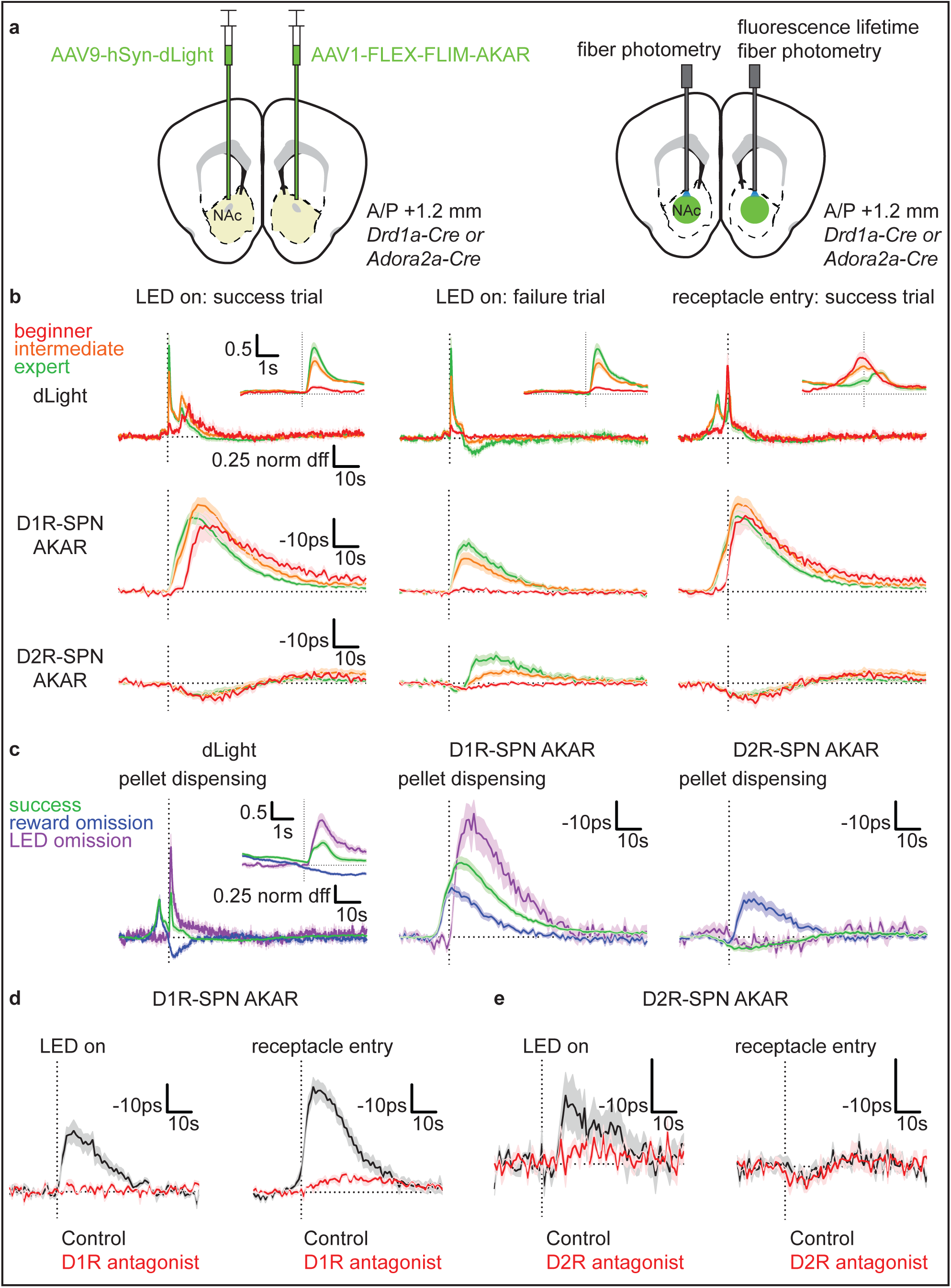
D1R- and D2R-SPN PKA activity are dynamically modulated by DA at different stages of learning. a) l*eft*, Schematic of a coronal section at 1.2 mm depicting injection of AAV9-hSyn-dLight and AAV1-FLEX-FLIM-AKAR into the NAc of *Drd1a-Cre* or *Adora2a-Cre* mice. *right*, Schematic of a coronal section depicting dual fiber photometry in which one fiber is used for intensity measurements of dLight fluorescence and the other for fluorescence lifetime measurements of FLIM-AKAR. The fibers were implanted 200μm above injection sites. b) dLight and FLIM-AKAR responses during learning. Top: average dLight responses (df/f normalized across all sessions for each mouse such that the 99 percentile response = 1) of *Drd1a-Cre* and *Adora2a-Cre* mice (n=16 mice consisting of 8 *Drd1a-Cre* and 8 *Adora2a-Cre* mice). Middle: average D1R-SPN FLIM-AKAR responses (delta lifetime) of *Drd1a-Cre* mice (n=14 mice consisting of 8 dLight+AKAR mice and 6 AKAR-only mice). Bottom: average D2R-SPN FLIM-AKAR responses (delta lifetime) of *Adora2a-Cre* mice (n=18 mice consisting of 8 dLight+AKAR mice, and 10 AKAR-only mice). Average responses are shown (line) with SEM (shaded area) aligned to the time (dashed vertical line) of “LED on” for success (*left*) and failure (*middle*) trials or “receptacle entry” for success trials (*right*). Beginner, intermediate, and expert trials are shown in red, orange, and green, respectively. Inset: magnified average dLight response. c) dLight and FLIM-AKAR responses in reward omission and rewarded LED omission trials aligned to the time of pellet dispensing (dashed vertical line). Average dLight (*left*), D1R-SPN AKAR (*middle*) and D2R-SPN AKAR (*right*) responses of expert mice on regular success (green), reward omission (blue), and rewarded LED omission trials (purple). n’s (mice) are as follows: dLight: 16, consisting of 8 *Drd1a-Cre* and 8 *Adora2a-Cre*, for success and reward omission trials and 8, consisting of 4 *Drd1a-Cre* and 4 *Adora2a-Cre*, for rewarded LED omission trials; D1R-SPN AKAR *Drd1a-Cre*: 14, consisting of 8 dLight+AKAR and 6 AKAR-only, for success and reward omission trials and 6, consisting 4 dLight+AKAR and 2 AKAR-only, for rewarded LED omission trials; D2R-SPN AKAR *Adora2a-Cre:* 18, consisting of 8 dLight+AKAR and 10 AKAR-only, for success and reward omission trials and 10 mice, consisting of 4 dLight+AKAR and 6 AKAR-only, for rewarded LED omission trials. Inset: magnified average dLight response. d) Effects of Type 1 dopamine receptor antagonism on behaviorally-induced transients in D1R-SPN PKA activity in *Drd1a-Cre* mice (n=4). Average responses of D1R-SPN FLIM-AKAR without (black) and with (red) drug aligned to the time of an LED cue previously associated with food delivery but with no food delivered (*left*, 5 trials/mouse) or to a free (i.e. no task related action required) reward aligned to the time of food pellet receptacle entry (*right*, 10 trials/mouse). Dopamine receptor antagonist was delivered IP at least 10 min before recordings began. e) As in panel (d) for D2R-SPN FLIM-AKAR in *Adora2a-Cre* (n=5 mice) without (black) and with (red) Type 2 dopamine receptor antagonism. *For all graphs, shaded area=SEM of averages across mice, total numbers of trials per each condition for b) and c) are given in the Extended Data Fig. 7.

In contrast, D2R-SPNs’ net PKA activity was minimally modulated in beginner animals (Fig. 3b and Extended Data Fig. 9). Furthermore, the small reduction in D2R-SPNs’ net PKA activity in success trials did not change across different stages of learning, suggesting that it is not caused by DA. On the other hand, failure trials of intermediate and expert animals, in which DA levels significantly drop below baseline, induced a significant increase in net PKA activity in D2R-SPNs (Fig. 3b). Similar increases in D2R-SPNs’ net PKA activity were seen in reward omission trials (Fig. 3c), consistent with this signal reflecting activation of PKA in these cells due to a rapid decrease in DA below baseline. We confirmed that the FLIM-AKAR modulation observed during behavior does not reflect movement artifacts by repeating lifetime measurements in separate cohorts of mice expressing mutant AKAR (FLIM-AKAR^T391A^) in D1R- or D2R-SPNs (Extended Data Fig. 8c).

These results reveal that PKA in D1R- and D2R-SPNs does not respond to the same features of DA dynamics and that PKA is not strongly modulated in the two cell types at the same time. Rather, D1R-SPN PKA is significantly activated even at early stages of learning by reward and continues to be activated by each reward-predicting cue and actual reward after learning whereas D2R-SPN PKA is only activated at late stages of learning by failure to achieve expected rewards.

With the exception of the small reduction in D2R-SPNs’ net PKA activity at the time of reward-predicting cue or reward, SPNs’ net PKA activity patterns could be explained by the observed dynamics of DA release that evolve across learning stages. To test a causal relationship between DA and SPN PKA modulation during behavior, we used the DA-receptor targeting pharmacology established above. Both the LED- and the reward-driven increase of D1R-SPNs’ net PKA activity were largely blocked by D1R antagonist (SKF 83566 hydrobromide, 3mg/kg IP) (Fig. 3d). Similarly, the reward omission driven activation of D2R-SPN PKA was blocked by D2R antagonist (Eticlopride hydrochloride, 0.5mg/kg IP) (Fig. 3e). This demonstrates that D2R-PKA activation during the behavior is mediated by D2Rs and that the basal tone of DA acting on D2Rs constitutively inhibits PKA allowing a dip in DA to transiently activate PKA in these cells. In contrast, the small reduction in D2R-SPNs’ net PKA activity after reward consumption was not blocked by D2R antagonist (Fig. 3e), consistent with a conclusion that this effect is independent of DA and D2Rs.

### Transient changes in DA neuron activity are sufficient to modulate SPN PKA activity

To establish the causal relationship between changes in VTA DA neuron firing, NAc DA levels and SPN PKA activity, we tested the sufficiency of manipulations DA neuron firing to modulate SPN PKA using optogenetics. We expressed activating (ChrimsonR^36^) or inactivating (stGtACR2^37^) opsins in DA neurons of (1) *DAT-IRES-Cre*, (2) *DAT-IRES-Cre; Drd1a-Cre*, and (3) *DAT-IRES-Cre; Adora2a-Cre* mice (Fig. 4a). In addition, we expressed dLight in the NAc of *DAT-IRES-Cre* mice and FLIM-AKAR in Cre-expressing SPNs in the double transgenic animals. We confirmed that we can bi-directionally change DA levels detected by dLight using the two opsins and that this modulation depends on the opsin stimulation protocol (Fig. 4b and e). Using this information, we calibrated Chrimson activation so that DA release from DA neuron optogenetic activation was matched to that of a natural food reward response (Fig. 4b). Under this stimulation strength, D1R-SPN net PKA activity strongly increased in a D1R-dependent manner (Fig 4c). A similar effect was achieved by stimulating DA neuron axons in NAc (Extended Data Fig. 10c).

**Fig. 4.**
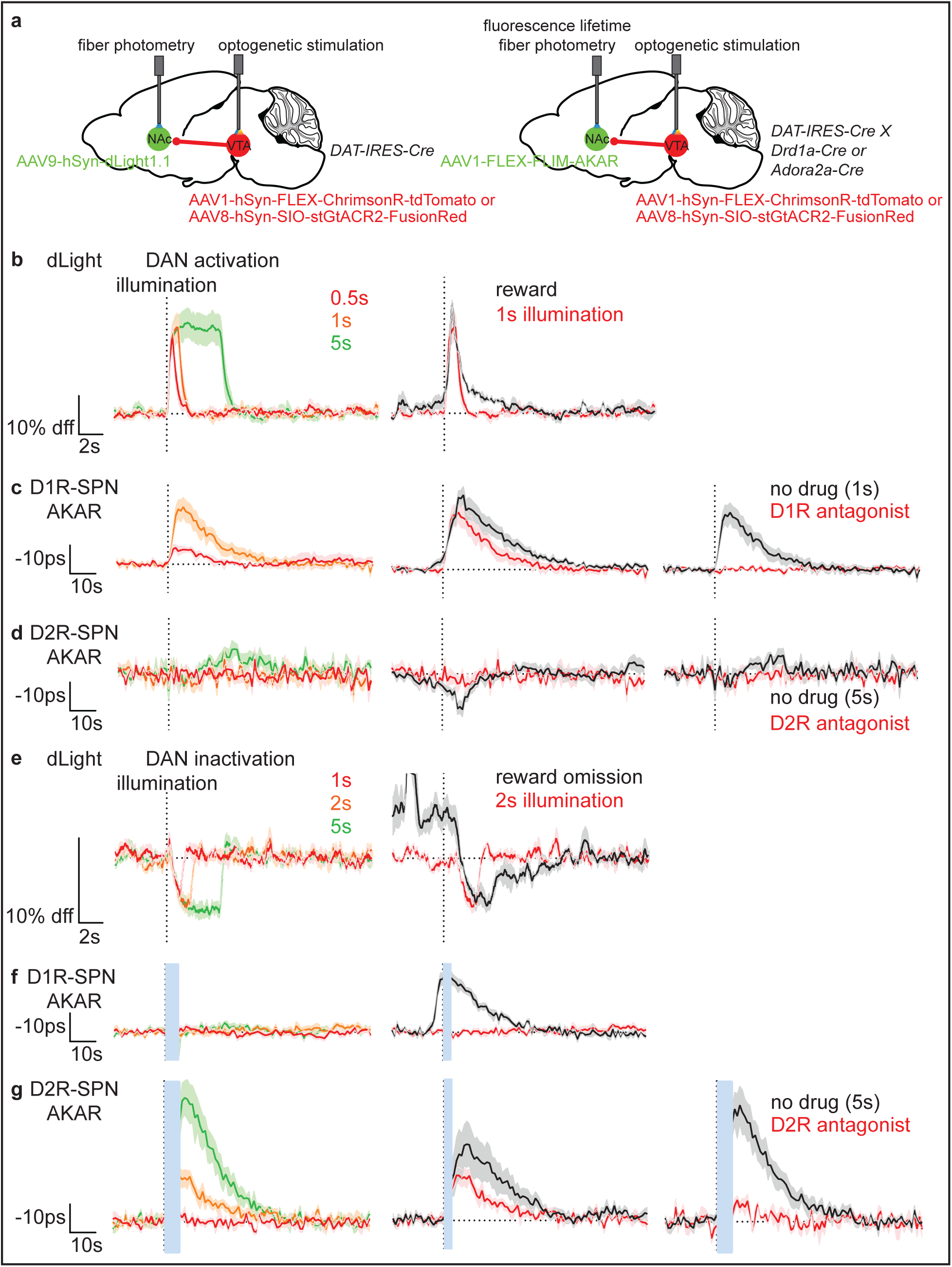
Transient changes in DA neuron activity are sufficient to modulate SPN PKA activity. a) Schematic of a sagittal section depicting viral expression and fiber implantation. *left*, AAV9-hSyn-dLight was injected into NAc of *DAT-IRES-Cre* mice. AAV1-hSyn-FLEX-ChrimsonR-tdTomato or AAV2-hSyn-SIO-stGtACR2-FusionRed was injected into VTA for DA neuron activation or inactivation, respectively. Fibers were implanted 200μm above the injection sites. *right*, As on the left except AAV1-FLEX-FLIM-AKAR was injected to NAc of *DAT-IRES-Cre; Drd1a-Cre* and *DAT-IRES-Cre; Adora2a-Cre* mice for, respectively, D1R-SPN and D2R-SPN FLIM-AKAR measurements. b) dLight response in *DAT-IRES-Cre* mice to optogenetic DA neuron activation. *left*, dLight responses (n=3 mice) to stimuli (20Hz train of 2ms pulses at 14.3mW) of different durations (red=0.5s, orange=1s, green=5s). *right*, dLight responses of naïve animals to food reward (black, n=3 mice) and the responses to 1s illumination (red, n=3 mice). Dashed line indicates the time of illumination onset for optogenetic trials. For food rewards, the response was aligned to its peak after receptacle entry and time-shifted for maximum overlap with an optogenetic response. c) As in panel (b) for D1R-SPN FLIM-AKAR responses in *DAT-IRES-cre; Drd1a-Cre* mice to 0.5 or 1 s optogenetic DA neuron activation (*left*, n=6 mice), food reward compared to 1s illumination (*middle*, n=6 mice), or 1s illumination without (black) or with (red) IP injection of a D1R antagonist at least 10min before recording (*right*, n=4 mice). d) As in panel (b) for D2R-SPN FLIM-AKAR responses in *DAT-IRES-Cre; Adora2a-Cre* mice (n=4 for each) to DA neuron activation with the exception that D2R antagonists were used for comparison to the 5s illumination response. e) dLight response of *DAT-IRES-cre* mice to DA neuron inactivation. *left*, dLight responses (n=4 mice) to stGtACR2 stimulation (continuous illumination, 5mW) to inhibit DA neuron firing for different durations (red=1s, orange=2s, green=5s). *right*, dLight responses of trained mice to reward omission (black, n=4 mice) and the responses to 2s illumination (red, n=4 mice). Dashed line indicates the time of illumination onset or expected reward delivery. The higher baseline dLight signal in reward omission response is due to the response to the LED cue that precedes reward omission (Fig. 2f). f) As in panel (e) for D1R-SPN FLIM-AKAR responses in *DAT-IRES-Cre; Drd1a-Cre* mice to DA neuron inactivation. *left*, D1R-SPN FLIM-AKAR responses to DA neuron inactivation of different durations (n=7 mice). *right*, D1R-SPN FLIM-AKAR responses of trained mice to the LED cue (~8s before the expected reward delivery) without food reward (black, n=7 mice) and the response to 2s illumination (red, n=7 mice). g) As in panel (e) for D2R-SPN FLIM-AKAR responses in *DAT-IRES-Cre; Adora2a-Cre* mice to DA neuron inactivation. *left*, D2R-SPN FLIM-AKAR responses to DA neuron inactivation of different durations (n=5 mice). *middle*, D2R-SPN FLIM-AKAR responses of trained mice to reward omission (black, n=5 mice) and the responses to 2s illumination (red, n=5 mice). *right*, Effect of D2R blockade on D2R-SPN FLIM-AKAR response to 5s DA neuron inactivation without (black, n=5 mice), and with (red, n=3 mice) IP injection of a D2R antagonist at least 10 min before recording. *For all graphs, shaded area=SEM of averages across mice. The average response of each mouse was calculated from 10 trials for optogenetic illumination and reward responses, and 5 trials for reward omission responses. Blue bars indicate the periods of blue laser illumination for stGtACR2 during which accurate FLIM-AKAR measurements were not possible. Reward responses of SPN FLIM-AKAR were acquired in separate cohorts of untrained animals. Reward-omission responses of dLight and SPN FLIM-AKAR were acquired from separate cohorts that were trained on the behavioral task.

In contrast, D2R-SPN net PKA activity was minimally modulated by DA neuron activation, suggesting that the small reduction in D2R-SPN net PKA activity induced by reward (Fig. 3b-e) is not DA dependent and that D2R-dependent inhibition of PKA is close to maximum at basal DA levels (Fig. 4d). To test if D2R-SPN PKA activity can respond to DA neuron activation at all, we activated DA neuron for 5s and 10s, which increase DA levels far beyond a natural food reward response (Fig. 4b and Extended Data Fig. 10b). This non-physiological stimulus resulted in a small but significant unexpected and delayed increase in D2R-SPN net PKA activity (Fig. 4d, Extended Data Fig. 10b), which did depend on D2Rs. Indirect circuit mechanisms, such as modulation of the activity of D2R-expressing cholinergic interneurons in the NAc, may have produced this paradoxical increase.

DA neuron inactivation had converse effects. D1R-SPN net PKA activity showed no significant change in response to DA neuron inactivation even at the longest inactivation protocol attempted (Fig. 4e-f). This suggests that the suppression of D1R-SPN net PKA activity by D1R antagonist (Fig. 1b) is most likely due to the inverse agonist effects of the drug and that there is a minimal effect of basal DA tone on D1R-SPN PKA activity. On the other hand, D2R-SPN net PKA activity was significantly increased in a D2R-dependent manner by a DA neuron inactivation protocol that decreased DA to a similar extent as reward omission in trained animals (Fig. 4e and g). These results are again consistent with a strong basal engagement of D2Rs by DA that allows rapid unbinding of DA from D2Rs to increase D2R-SPN PKA activity. In summary, optogenetic manipulation of DA neurons re-capitulates the asynchronous modulation of D1R- and D2R-SPN PKA activity by DA increases and decreases, respectively, observed during the behavioral task.

### Selective PKA inhibition in SPNs partially impairs learning

Given the separate modulation of D1R- and D2R-SPN PKA activity during learning, we investigated if a selective inhibition of PKA activity in each neuron class differentially affects learning. To achieve cell-type specific inhibition of PKA activity, we virally overexpressed PKI_α_, an endogenous inhibitory peptide for PKA, which, we previously showed, effectively blocks PKA activation *in vitro*^31^ and *in vivo*^32^. We injected AAV1-FLEX-PKIalpha-IRES-nls-mRuby2 or AAV1-Cag-FLEX-eGFP (control) into *Drd1a-Cre* and *Adora2a-Cre* mice to selectively inhibit PKA in D1R- and D2R-SPNs, respectively (Fig. 5a). 10-14 days after injection, the mice were trained on the behavioral task for 11 days, followed by 7 days of extinction sessions in which no food reward was provided during the task while the mice remained food restricted.

**Fig. 5.**
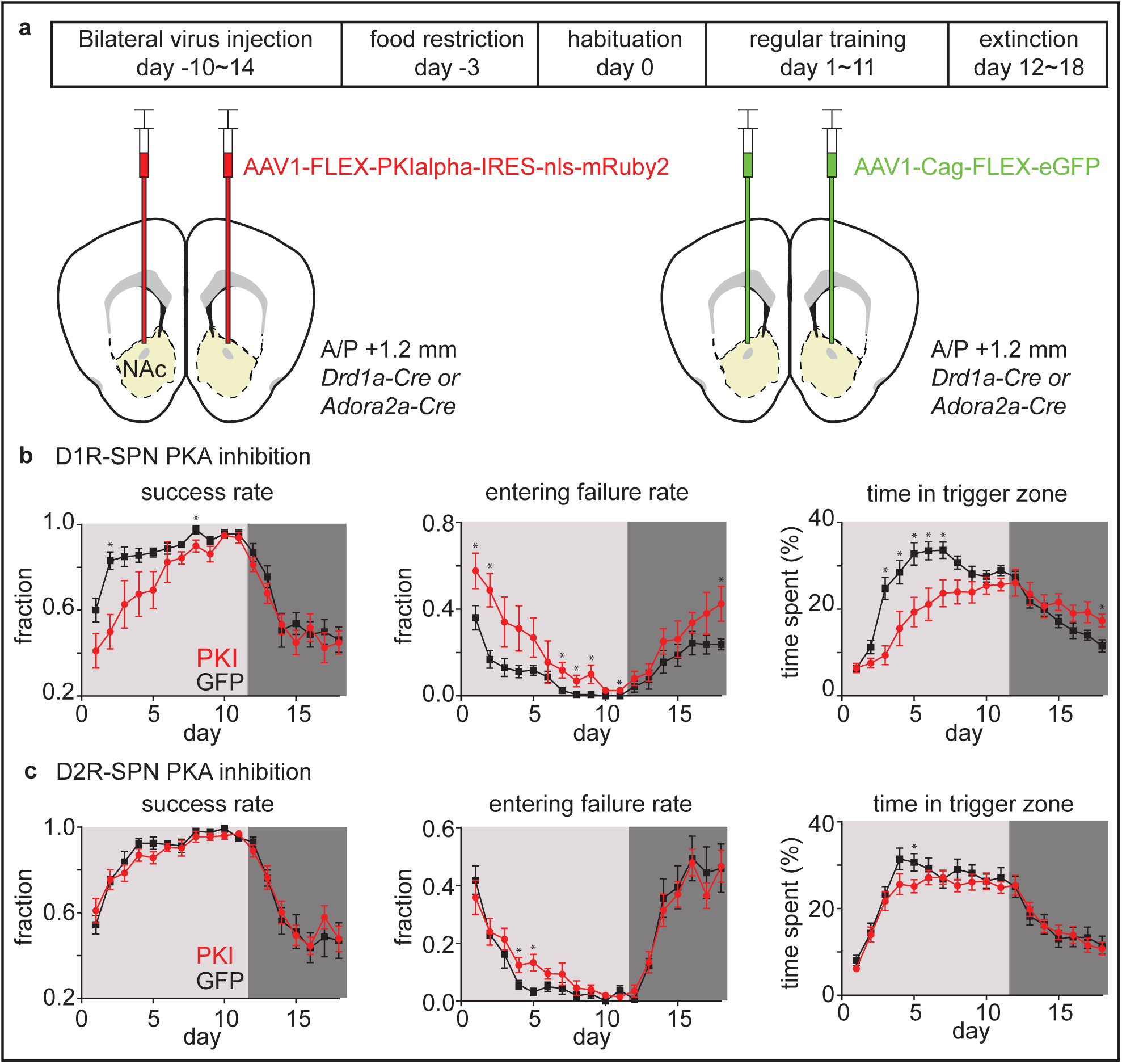
Selective PKA inhibition in SPNs partially impairs learning. a) Schematic describing a strategy to investigate the effect of D1R-SPN or D2R-SPN PKA inhibition on behavior. AAV1-FLEX-PKIalpha-IRES-nls-mRuby2 was injected into NAc of *Drd1a-Cre* or *Adora2a-Cre* mice to selectively inhibit PKA in D1R-SPN or D2R-SPN, respectively. For control groups, AAV1-Cag-FLEX-eGFP was injected into NAc, instead. 10-14 days after surgery, mice were started on a behavior schedule that includes 1 day of habituation (day 0), 11 days of regular training (day 1~11), and 7 days of extinction (day 12~18). b) Effect of D1R-SPN PKA inhibition on learning. *left*, Success rate (number of rewarded trials/total number of trials). *middle*, Entering failure rate (number of receptacle zone entering failure trials/total number of trials). *right*, Time spent in the trigger zone (time spent in a zone/total session time). The averages of PKI (n=8 mice) and GFP (n=8 mice) groups are shown in red and black, respectively. For all graphs, error bar=SEM of averages across mice, light gray=days of regular training, dark gray=days of extinction. * represents a p-value < 0.05 for Welch’s t-tests. c) As in panel b) for the effect of D2R-SPN PKA inhibition on learning. n=10 mice (PKI) and 8 mice (GFP).

As predicted from a large activation of D1R-SPN PKA at the time point of DA release (Fig. 3b), PKI expression in D1R-SPNs significantly slowed learning as measured by a success rate (Fig. 5b). The lower success rate was mostly due to the higher rate of entering failure (failure of entering the receptacle zone after the LED cue on time) (Fig. 5b). Although average movement speed of the PKI group was lower than that of the GFP group (Extended Data Fig. 11a), this effect cannot fully explain slower learning because, 1) PKI mice were physically capable of moving sufficiently fast to enter the receptacle zone (the plateau success rates were not significantly different between the two groups), and 2) the time spent in the LED trigger zone also increased more slowly for the PKI group, which cannot be explained by PKI mice moving more slowly or simply staying in the corners of the box (Fig. 5b). Interestingly, there was also a slight increase in entering failure rate in PKI group during extinction trials (Fig. 5b), suggesting that D1R-SPN PKA activation by a reward associated cue (Fig. 3b and c) may participate in sustaining a learned behavior in the face of omission of rewards.

Consistent with lack of strong D2R-SPN PKA modulation in the early phase of learning (Fig. 3b), PKI expression in D2R-SPNs did not change the learning rate in the first few days (Fig. 5c). In line with the activation of D2R-SPN PKA in the failure trials in intermediate and expert animals (Fig. 3b), there was a slight slowing of the learning rate in the late phase of learning (day 4~5) in PKI group represented by higher entering failure rate and shorter time spent in the LED trigger zone (Fig. 5c). In contrast to D1R-SPN PKI expression, there was a minimal effect of D2R-SPN PKI expression on the overall movement (Extended Data Fig. 11b). Surprisingly, there was no significant effect of D2R-SPN PKA inhibition on the extinction trials (Fig. 5c), which suggests that D2R-SPN PKA activation in the NAc caused by reward omission (Fig. 3c) is not required for extinction. Overall, these PKI experiments demonstrate that D1R- and D2R-SPN PKA participate in different stages of reward based learning, consistent with the asynchronous activation of D1R- and D2R-SPN PKA during learning.

## Discussion

Using multi-channel fiber photometry and fluorescence lifetime photometry (FLiP), we measured transients in VTA DA neuron activity, NAc DA levels, and net PKA activity in D1R- or D2R-SPNs in freely-moving animals learning an operant conditioning task. We found robust RPE-encoding in the activity of VTA DA neurons, as measured either at the cell body or the axons in NAc, and in NAc DA levels. Furthermore, the RPE-encoding bidirectional transients in DA levels were sufficient to explain dynamic PKA signaling in each class of neuron. D1R-SPN net PKA activity increased in response to rewards and reward-predicting cues throughout and after learning whenever there was a phasic DA increase. In contrast, D2R-SPN net PKA activity was only modulated by dips in DA levels that appeared after learning and were caused by performance errors or reward omission. Using pharmacological and optogenetic manipulations, we demonstrated that behaviorally-evoked DA transients, along with differential basal modulation of PKA by D1Rs and D2Rs, can directly cause the modulation of SPN PKA activity and explain the asynchronous patterns of SPN PKA modulation in the two cell classes. Lastly, the temporally-dissociated modulation of SPN PKA activity by DA is consistent with the differential effects of cell-type specific inhibition of PKA, which revealed that only PKA inhibition in D1R-SPNs significantly reduces the initial learning rate.

### Relationships between DA neuron activity and DA release

The relationship between VTA DA neuron action potential firing and NAc DA release has been put into question by a recent study showing that phasic VTA DA neuron firing does not fully account for NAc DA release patterns in an expert animal performing a freely moving behavior task^17^. This study suggested that the dissociation between DA neuron soma activity and DA release might be explained by a significant modulation of DA release locally at DA neuron terminals, such as a modulation by cholinergic interneurons^3839^^40^. Here, using multi-channel fiber photometry to monitor population activity of VTA DA neurons and DA release simultaneously, we found that VTA DA neuron activity (measured by Ca^2+^ level) and DA release maintain a strong moment-to-moment correlation across all stages of learning (Fig. 2e and Extended Data Fig. 4b). Furthermore, we observed that reward history affects both VTA DA neuron activity and DA release during the baseline period (no cue or reward during this period) unlike the result of the previous study^17^ in which the history only affected DA release not VTA DA neuron firing rate during this period (Extended Data Fig. 6).

These conflicting results may arise from a difference in how we measure VTA DA neuron activity. Our study used a genetically encoded Ca^2+^ indicator (GECI) whereas Mohebi et al. performed *in vivo* electrophysiology recording. Although a GECI has a limited capacity to detect small changes in Ca^2+^ level without action potentials^41^, jRCaMP1b could have reported a global but subtle change in Ca^2+^ level that does not result in action potential firing in DA neurons during the baseline period in our study. Another possibility is a difference in a trial structure, as individual trials were separated by an inter trial interval (ITI) of >=120s and 5-10s, in our study and Mohebi et al., respectively. The behavior task with a shorter ITI may have engaged an additional circuit, such as cholinergic interneurons in NAc, which could dissociate DA release from DA neuron soma activity. Lastly, the difference could partly come from a slight difference in recording sites for VTA given different animal models (rats vs. mice) were used. The heterogeneity in information encoded by DA neurons in different regions of VTA^2^ supports this possibility. Thus, although our findings do not negate the role of local modulation of DA release, they support a model that RPE-encoding transients in VTA DA neuron somatic activity effectively control DA levels in the NAc during learning.

### DA or DA neuron activity as RPE

Many studies have demonstrated that both VTA DA neuron activity and DA release in the NAc are modulated in patterns that reflect RPE^11,12,13,14,15,16,17^. However, most examined either animals just beginning to learn a task or highly-trained animals. Due to the late appearance of RPE-like modulation of DA neuron firing during training in many of these studies, it has been suggested that RPE-like DA transients may not be correlated with adaptive changes in behavior and cannot provide a behaviorally relevant RPE teaching signal^9^. Few studies^15^ have measured DA neuron activity and DA release in freely-moving animals during training. Here, by recording neural signals in individual animals in every session across learning, we demonstrate that both DA neuron activity and DA release patterns in the NAc show a progressive change that follows RPE (Fig. 2 and 3) and are correlated with behavioral performance throughout learning (Extended Data Fig. 3c).

Other studies have suggested that DA levels may encode or generate movements^9,42,43,44^. Particularly in dorsal striatum, which receives DA input predominantly from the substantia nigra pars compacta (SNc), DA levels are correlated with and appear causally related to movement onset. Indeed, in both VTA and NAc, we find a weak correlation between DA neuron activity/DA release and movement (Extended Data Fig 5); however, both signals were better explained by reward-predicting cues and rewards. These cue and reward driven DA signals may be, in part, encode cue- and reward-evoked movements, yet they encode the features of RPE, regardless. Furthermore, our result is consistent with the previous findings^42,44,45^ that DA signals in dorsal and ventral striatum preferentially (but not exclusively) encode movement and reward, respectively.

Coddington et al.^9^ further proposed a model in which both SNc and VTA DA neurons encode the timing of action. In this model, RPE-like patterns of DA neuron activity result from the summation of sensory cue-related and movement initiation-related components, and are seen only in well-trained animals. In contrast, we observed a gradual change in both DA neuron activity and DA release patterns consistent with RPE across different stages of learning. The differences between our results and those of the previous study^9^ may arise from experimental differences, such as whether an animal is head-restrained or freely-moving and whether the task is Pavlovian (classical) or instrumental. On the other hand, gradual changes in cue and reward response of DA neurons were not perfectly synchronized in that the reward response continues to decrease in the late stage learning even when the cue response increase has plateaued (Extended Data Fig. 3c), which is consistent with the previous study^9^. This suggests different mechanisms maybe involved in a computation that results in increase in cue response and reduction in reward response of DA neurons.

Due to the experimental limitation of having a low number of trials per session, we were not able to formally test the relationship between the value of expected reward and DA signals proposed by the previous studies^16,17^. Yet, consistent with these studies, we observed the effects of trial history on DA levels in the NAc (Extended Data Fig. 6) within a single session, which may allow DA level to encode a reward rate or a value in addition to an RPE teaching signal.

In summary, our results do not disprove the additional roles of DA proposed by the previous studies, but, rather, strengthen a classical reinforcement model, where RPE encoding DA signal acts as a useful teaching signal to downstream SPNs.

### SPN PKA modulation and function during behavior

Direct measurements of the downstream effects of RPE-encoding DA transients on SPNs have been difficult to achieve due the challenges of monitoring intracellular signaling in behaving animals (although see Goto et al.^46^, Yamaguchi et al.^47^, and Ma et al.^48^). Employing the fluorescence lifetime photometry technique in combination of pharmacological and optogenetic manipulations, we successfully monitored the dynamics of net PKA activity in SPNs during behavior. We showed that PKA activity in D1R- and D2R-SPNs follow the patterns of DA release and established the causal relationship between the two. These indicate that, despite the potential contributions of many GPCRs and Ca^2+^-dependent adenylyl cyclases, DA, at times of its phasic modulation, has a strong control over PKA in both D1R- and D2R-SPNs. However, there was an exception to this relationship in that there was a small reduction in D2R-SPN net PKA activity after reward consumption. This small change in net PKA activity appear to be DA and D2R independent given that this response was not blocked by D2R antagonist (Fig. 3e) and could not be replicated with optogenetic DA neuron activation (Fig. 4d).

Importantly, we observed context and cell-type dependent regulation of intracellular pathways by DA: D1R-SPN net PKA activity was enhanced by DA increases in all, including the earliest, phases of learning whereas D2R-SPN net PKA activity was modulated significantly only by DA decreases that emerge during the late phases of learning and are triggered when expected rewards are not delivered. Furthermore, pharmacological and optogenetic perturbations revealed the origin of this difference by providing direct *in vivo* evidence supporting the classical hypothesis that, due to differences in their ligand binding affinity, D2Rs but not D1Rs are occupied by basal DA levels^29,30,49^. Additionally, the activation of PKA in D2R-SPNs following transient dips in DA indicates that basal DA provides continuous inhibition of PKA *in vivo*, suggesting that basally active DA-bound D2Rs are not desensitized (or internalized) and continually suppress adenylyl cyclases.

This asynchronous engagement of D1R-SPN and D2R-SPN PKA activity had a functional implication for basal ganglia dependent learning. Selective inhibition of PKA activity in D1R-SPN using the genetically-encoded PKI altered learning starting from its initial phases whereas a mild effect of D2R-SPN PKA inhibition was observed only at the late phase of learning (Fig. 5). This is consistent with the fact that D1R-SPN PKA is strongly modulated starting in the beginning of training whereas D2R-SPN PKA is significantly activated only during the late phase of training.

Interestingly, we did not observe a complete impairment of learning when we inhibited PKA activity in D1R- or D2R-SPNs. This could be due to 1) incomplete blockade of PKA activity in either classes of SPNs 2) PKA independent cellular plasticity and/or 3) other brain regions involved in learning that may compensate for the function of NAc. Possibility 1 is less likely given we have previously observed a near complete blockade in PKA activation by PKI in both *in vitro*^31^ and *in vivo*^32^. In addition, D2R-SPN PKA was strongly activated during reward omission trials, but PKA inhibition in D2R-SPNs did not have a significant effect on the extinction sessions. It is possible that different parts of the striatum may be involved in extinction^50^. Despite these caveats, our results demonstrate that DA dependent modulation of PKA in SPNs is at least partially involved in reinforcement learning.

### Implications for models of basal ganglia and reinforcement learning

In summary, our findings are not consistent with a model in which bidirectional dynamics of DA levels simultaneously modulate PKA activity in D1R- and D2R-SPNs. Rather, we propose a dichotomous model (Extended Data Fig. 12) of DA and basal ganglia dependent learning in which PKA activities in D1R- and D2R-SPNs are asynchronously engaged to induce plasticity in each cell type, mediating the action of RPE encoding DA signals during different phases of learning. More specifically, we propose that the direct (striatomesencephalic) and indirect (striatopallidal) pathways have independent functions during learning: the former forming an initial association between an action and an outcome and the latter refining the learned association.

It is still mysterious how the slow kinetics of net PKA activity allow temporally precise associations of past actions and action outcomes. The slow kinetics observed by FLIM-AKAR are not a result of saturating PKA or phosphatase capacity by the sensor^32^, and thus provide a good estimate of the kinetics of net phosphorylation of diffusible PKA substrates in SPNs. It is unlikely that there is a large difference in the phosphorylation kinetics of anchored (e.g. synaptic) vs. diffusible PKA substrates^51^. Therefore, our findings support a model, in which the activation of a second parallel pathway (such as CaMKII) preceding PKA activation provides temporal precision and is required for synaptic plasticity during learning^24,52^.

## Acknowledgement

This work was supported by NIH (NINDS R35NS105107, B.L.S.), Howard Hughes Medical Institute (B.L.S.), and the Sackler Scholar Programme in Psychology (S.L). Graphical illustration was provided by Sigrid Knemeyer.

## Author contributions

S.L. designed the study, developed the photometry system, collected and analyzed data, and wrote the manuscript. B.L. collected and analyzed data, and edited the manuscript. Y.C. conceived the study and edited the manuscript. T.P. and L.T. developed the dLight sensor and shared expertise. B.L.S. designed and supervised the study and wrote the manuscript.

## Methods

### Animals

Experimental manipulations were performed in accordance with protocols approved by the Harvard Standing Committee on Animal Care following guidelines described in the US National Institutes of Health Guide for the Care and Use of Laboratory Animals. *Drd1a-Cre* (B6.FVB(Cg)-Tg(Drd1-cre)EY262Gsat/Mmucd, 030989-UCD) and *Adora2a-Cre* (B6.FVB(Cg)-Tg(Adora2a-cre)KG139Gsat/Mmucd, 036158-UCD) mice on *C57BL/6J* backgrounds were acquired from MMRRC UC Davis. *DAT-IRES-Cre* (*B6.SJL-Slc6a3tm1.1(cre)Bkmn/J*, 006660) and *C57BL/6J* (000664) mice were acquired from the Jackson Laboratory. *DAT-IRES-Cre; Drd1a-Cre* and *DAT-IRES-Cre; Adora2a-Cre* mice were bred in house by crossing heterozygous parent lines. All transgenic animals used for experiments were heterozygous for the relevant Cre allele.

### Viruses

Recombinant adeno-associated viruses (AAVs of serotype 1, 8, 9) were used to express transgenes of interest in either Cre-recombinase dependent or independent manner. AAVs were packaged by commercial vector core facilities (Addgene, Boston Children’s Hospital Vector Core, Janelia Vector Core, Penn Vector Core, UNC Vector Core) and stored at −80°C upon arrival. Viruses were used at a working concentration of 10^12^ to 10^14^ genomic copies per ml. 300 nl of virus was used for all experiments except for PKI experiments where we used 600 nl of virus at 1.2 × 10^14^ genomic copies per ml and 300 nl of virus at 4 × 10^13^ genomic copies per ml for *Drd1a-Cre* and *Adora2a-Cre*, respectively (we used a lower volume and titer for the latter due to a toxicity issue).

### Surgery

Inhaled isoflurane was used as anesthesia. Virus was stereotactically injected into either NAc core (anteroposterior (AP) +1.2 mm, medio-lateral (ML) +/−1.3 mm relative to bregma; dorsoventral (DV) 4.1mm below brain surface) or VTA (AP −3.3 mm, ML +0.48 mm, DV 4.5 mm). For fiber photometry or optogenetic experiments, an optical fiber (MFC_200/230-0.37_4.5mm_MF1.25_FLT mono fiber optic cannula, Doric Lenses) was implanted 200μm above the injection site. For intraventricular injections, an infusion cannula (C313GA/SPC GUIDE ACUTE 22GA, Plastics One) was implanted into the ventricle (AP +0.5 mm, ML −1.75 mm, DV 1.5 mm) of the hemisphere opposite to the fiber implantation site at an angle (20 degrees from the axis of stereotactic holder).

### Behavior

To motivate animals to perform the visual cue guided operant task, we food restricted mice such that they remained at 80~90% of their initial weight. Mice were given 2~3 g of regular chow daily in addition to the variable number of 20 mg dustless precision chocolate flavor pellets (F05301, Bio Serv) consumed during the task. Food restriction was started at least one day before commencing the behavior experiments. During the task, a mouse (sometimes connected to a patch cord) was allowed to freely move inside an 8×16 inch behavior box, which contained a pellet receptacle and a white LED on one of the short walls. Receptacle entry was detected by an infrared sensor installed inside the receptacle. Animal movements were captured by cameras (WV-CP504, Panasonic or FL3-U3-13E4M, PointGrey) connected to Ethovision or Bonsai software that controlled all behavioral apparatuses and were synchronized with MATLAB software used to acquire photometry data.

The behavioral task structure is the following. After the enforced 120 s ITI, the mouse can self-initiate a trial by entering the trigger zone (a zone opposite to the wall containing the pellet receptacle indicated in green in Fig. 2a but not marked in the actual behavior box) and staying in the zone for 3 consecutive seconds. If the mouse enters the trigger zone during the enforced ITI or exits the trigger zone before 3 s after entering the zone, the trial is not initiated. If the mouse succeeds in initiating the trial, the LED above the receptacle turns on, signaling the start of the trial. Once the LED turns on, the mouse must enter the LED zone (a zone near the wall containing the pellet receptacle indicated in blue in Fig. 2a but not marked in the box) with in 5 s in order to continue with the trial. If the mouse fails to enter the LED zone within 5 s, the trial gets terminated, and the LED turns off. If the mouse enters the LED zone in time, it has to stay in the LED zone for additional 3 s. After this enforced waiting period, a single 20 mg pellet is dispensed, and the LED turns off. If the mouse prematurely exits the LED zone during the waiting period, the trial is terminated, and the LED turns off. After termination of the trial either by a success or a failure, the next enforced ITI of 120 s starts. Mice were trained for 11 days starting after 1 day of consisting of a 40-minute habituation session during which they were given 10 free pellets in the receptacle. On day 12, the reward was omitted from 25% of successful trials to collect “reward omission” data. On day 13, the LED was turned off for the entire session to collect “LED omission data”. Most mice reached 90% success rate (# of success trials/# of total trials) in 9 days (Extended Data Fig 3a.).

Each behavior session was run until a mouse initiated 20 trials (including both success and failure trials) or a 2 hr time limit was reached except reward omission days (day 12) when the session continued until 5 reward omission trials were collected. Because mice had to self-initiate trials, session durations were variable and between 50 min ~2 hr. Most sessions included 20 trials except the first few sessions at the start of training and the extinction sessions of PKI experiments (9 out of 806 sessions for photometry experiments, 15 out of 611 sessions for PKI experiments).

To compare signals across different stages of learning across all animals, we defined a “beginner” stage as days in which the success rate was below 50% of the maximum (plateau) success rate, an “intermediate” stage as days in which the success rate was between 50~90% of the maximum success rate, and an “expert” stage as days in which the success rate was above 90% of the maximum success rate.

### Fluorescence lifetime photometry (FLiP)

FLiP was carried out with the optical system described in Extended Data Fig. 1 and using the method described in detail in Lee et al. (2019). Briefly. All filters in the system were purchased from Semrock. A pulsed laser (BDS-473-SM-FBE Becker and Hickl – BH – operating at 50Mhz) was used as the light source to excite FLIM-AKAR. For fluorescence detection, a high-speed hybrid photomultiplier tube (PMT, HPM-100-07-Cooled, BH) controlled by DCC-100-PCI (BH) was used. The hybrid PMT was connected to an SPC-830 (BH), a time correlated single photon counting (TCSPC) board, which detects the time delay between the pulsed excitation and the photon detection by the PMT. The data was collected by custom software written in MATLAB, which calculated the average lifetime of detected photons at 1s intervals. This interval for average lifetime measurements was empirically determined to have enough photons to accurately estimate the lifetime (>200,000 photons/measurement) without running into a photon count limit (~1,000,000 photons/s) of the TCSPC board. The typical excitation power needed to generate the appropriate rate of photons (~400 kHz) for TCSPC was 0.6~1 µW (measured at the output end of the patch cord).

To estimate the change in lifetime of FLIM-AKAR, we calculated the average lifetime for each 1s time bin by measuring the mean photon arrival time (the population mean of the delay between the pulsed excitation and the fluorescence photon arrival as described) using the following equation^53^:

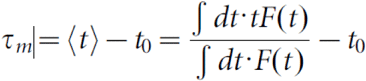

in which F(t) is the photon count of a fluorescence lifetime decay curve at time bin t, and t_0_ is the offset of the lifetime histogram, which can be estimated by fitting a double exponential curve to a lifetime histogram. We performed this calculation for the 0-8 ns time range in a lifetime histogram as this interval was minimally contaminated by a secondary fluorescence peak resulting from the autofluorescence of a fiber. The length of the patch cord was chosen to maximize the time separation between the sensor fluorescence peak and the fiber autofluorescence peak (~10 ns time delay for light to travel from the one end of the patch cord to the other end and back). The lifetime of FLIM-AKAR was reported as a change in lifetime (delta lifetime), which was calculated by subtracting the average lifetime of a baseline period (a period before the event of interest) from the average lifetime transient.

### Fiber photometry and optogenetics

Both fiber photometry (fluorescence intensity fiber photometry) and optogenetics were carried out with the optical system described in Extended Data Fig. 1. All filters in the system were purchased from Semrock except for a Doric Minicube (FMC5_E1(465-480)_F1(500-540)_E2(555-570)_F2(580-680)_S, Doric Lenses) that houses multiple dichroic mirrors and filters.

For fiber photometry, 470nm (M470F3, Thorlabs) and 565nm (M565F3, Thorlabs) LEDs were used to excite dLight and jRCaMP, respectively. Both LEDs were frequency modulated by a digital signal processor system (RX8-5-12, Tucker-Davis Technology) to carry out locked in amplification of PMT outputs. The average power levels of LEDs (measured at the output end of the patch cord) were 9.3 µW, 5.4 µW, and 16.2 µW for dLight, soma jRCaMP, and terminal jRCaMP excitation. For fluorescence detection, H7422-40 (Hamamatsu) and H10770(P)A-40 (Hamamatsu) PMTs were used. PMTs were connected to a low noise current preamplifier (SR570, Stanford Research Systems). The signal generated from SR570 was locked-in-amplified by RX8-5-12 using the frequency of the LED used to excite the sensor that fluoresces in the light spectrum assigned to a corresponding PMT. The locked-in-amplified signal was collected by a data acquisition board (PCI-6115, National Instruments), which was controlled by the custom software written in MATLAB. The raw fluorescence data was collected at 1 kHz. It was subsequently smoothed by a moving average filter (width of 200 ms) and down-sampled to 100 Hz. The relative change in fluoresce df(f)/f_0_ = (f(t)-f_0_)/f_0_ was calculated using f_0_ equal to the average of the baseline period (20 s before the trigger zone entry). For comparisons and averaging across mice, df/f of an individual mouse was normalized to the maximum df/f value across all sessions for the mouse.

For activation of Chrimson, a 593.5nm laser (SKU: YL-593-00100-CWM-SD-03-LED-F, Optoengine) was used with 2 ms laser pulses delivered at 20Hz. Reported laser powers were measured by a digital optical power meter (PM100D, Thorlabs) at the end of the patch cable, while the laser was operating in a continuous mode. For activation of stGtACR2, a 473nm laser (MBL-III-473, Optoengine) was used. For all stGtACR2 activation experiments, the laser was operated in a continuous mode (without pulsing). Reported laser powers were measured in the same manner as for the Chrimson experiments.

### Pharmacology

The following concentrations of drugs were used for intraperitoneal (IP) injection: D1R agonist (SKF 81297 hydrobromide, 10 mg/kg), D1R antagonist (SKF 83566 hydrobromide, 3 mg/kg), D2R agonist (Sumanirole maleate, 4 mg/kg), D2R antagonist (Eticlopride hydrochloride, 0.5 mg/kg), D2R antagonist (Sulpiride, 20 mg/kg), A2AR agonist (CGS 21680 hydrochloride, 4 mg/kg), and muscarinic receptor antagonist (Scopolamine hydrobromide, 10 mg/kg). Eticlopride hydrochloride was used for all D2R antagonist experiments unless specified otherwise. All drugs were dissolved in sterile saline except Sulpiride and CGS 21680 hydrochloride, which were dissolved in DMSO/saline mixture (10% v/v). Drug solutions were IP injected in 0.1 ml solution/10 g of mouse using a fine needle insulin syringe (BD 324911, BD Bioscience). On average, it took 30~60 s from the beginning of scruffing a mouse to the end of IP injection. For intraventricular (IV) injection, Sulpiride (2 mg/ml) and CGS 21680 hydrochloride (0.4 mg/ml) dissolved in DMSO/saline mixture (10% v/v) were used. Drug solutions were injected in a fixed volume of 2.5 µl at a speed of 250 nl/min by a high-precision syringe pump (70-3007, Harvard Apparatus) that was connected to an infusion cannula (C313GA/SPC GUIDE ACUTE 22GA, Plastics One).

### Deconvolution of fluorescence intensity photometry data

To gain a better understanding of the underlying neural activity generates the measured fluorescence transients, we deconvolved fluorescence signals into population events using the one-dimensional constrained deconvolution algorithm developed by Pnevmatikakis et al. (2016), which assumes that the sensor fluorescence follows an autoregressive process. Before deconvolution, we smoothed the raw fluorescence data collected at 1 kHz by a moving average filter (200 ms width) and down-sampled the data to 10 Hz. We subsequently calculated df/f using the bottom 5^th^ percentile value of a rolling time window of 300 s as f_0_. This allowed us to perform deconvolution on fluorescence signal for an entire session without introducing a trial structure to the data. The deconvolution of df/f was performed using a conic programing method and a second order autoregressive process. Correlations between signals were calculated by computing a correlation between the number of population events in each 1s time interval.

### Encoding model

To more objectively reveal the relationships between behavior and fluorescence signals, we created a generalized linear model to predict the deconvolved fluorescence signal (dLight and jRCaMP) from observed stimuli and behavioral parameters. We processed both observed stimuli and behavioral parameters into time bin of 100 ms to match the time bin of the deconvolved fluorescence signal. We constructed additional explanatory variables by introducing multiple time shifts to the observed stimulus and behavioral parameter variables. For continuous variables (speed, acceleration, rotation, position), −0.3 ~ +0.3s time shifts were introduced; for event variables (movement initiation, cue, reward delivery, receptacle entry), −0.3 ~ +1.0s time shifts were introduced. We reorganized the data from a session into a trial structure in order to have a fixed distribution of the number of trials when we split the data into fitting and testing sets. Model parameters (coefficients) were learned from fitting sets and evaluated using testing sets (5 fold cross validation). We performed Lasso on the data from all sessions of each learning stage and each mouse to find a minimum set of explanatory variables that allows the mean squared error (MSE) to be within 1 standard error from the minimum MSE (a standard error of MSE was calculated from 5 cross validation sets). We decided on the final common set of explanatory variables by selecting all variables selected by Lasso across different learning stages. With the final set of explanatory variables, we performed a linear regression. The contribution of each variable category (kinematics, movement initiation, position, cue, reward, accuracy, previous trial) was calculated by setting the coefficients of all the variables assigned to each category as zero and computing the correlation between the actual fluorescence signal and the signal predicted by the model: contribution = (R^2^_full_ – R^2^_partial_) / R^2^_full_.

---

**Extended Data Fig. S1.**
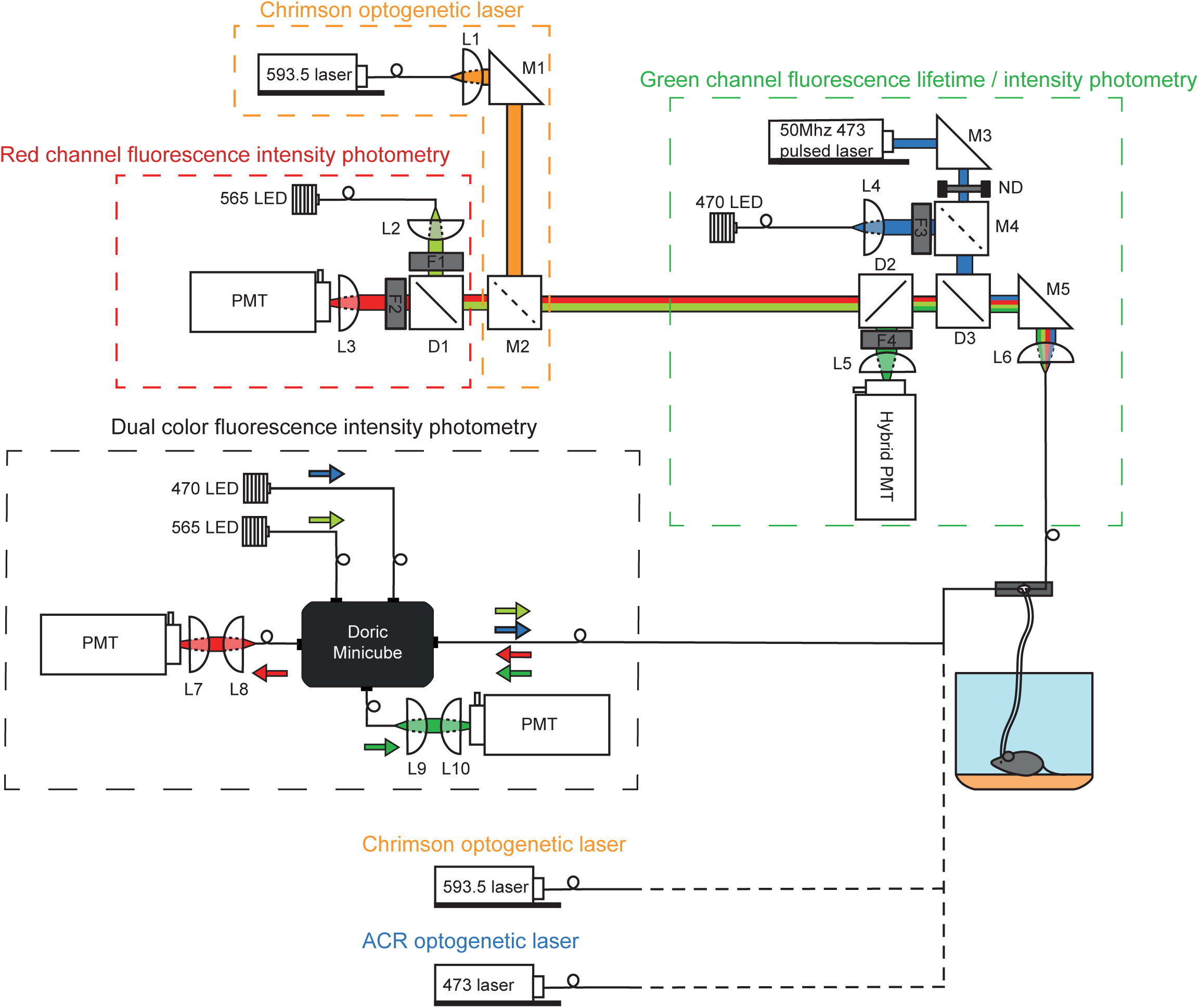
Multi-purpose photometry system for FLiP, fiber photometry, and optogenetics. The system consists of two independent multi-color photometry units. The top photometry unit consists of three sub-components used for: (1) red channel fluorescence photometry, (2) Chrimson optogenetic laser activation, and (3) green channel fluorescence lifetime and intensity photometry. For (1), red channel photometry was accomplished using a fiber coupled 565nm LED (M565F3, Thorlabs) for excitation whose output was collimated in free-space by L2 and filtered by F1 (554/23, Semrock). Red fluorescence was separated from the excitation light by dichroic D1 (573LP, Semrock), filtered by F2 (630/60, Semrock), and focused onto a PMT (H10770(P)A-40, Hamamatsu) by L3. For (2), Chrimson optogenetic light was provided by a fiber coupled 593.5nm laser (SKU: YL-593-00100-CWM-SD-03-LED-F, Optoengine) whose output was collimated by L1 and combined with the red photometry path via M2, a mirror that can be inserted or removed, respectively, for Chrimson optogenetic stimulation or red channel photometry. For (3)’s green channel fluorescence lifetime measurement mode, a 50Mhz 473nm pulsed laser (BDS-473-SM-FBE, Becker & Hickl) was fed through a rotating neutral density filter for power adjustment, reflected by D3 (488LP dichroic, Semrock), and focused onto a patch cable by L6. Emission light was passed through D3, reflected by D2 (532LP dichroic, Semrock), filtered by F4 (517/22, Semrock), and focused by L5 to a high-speed hybrid PMT (HPM-100-07-Cooled, Becker and Hickl). The hybrid PMT was connected to a time correlated single photon counting board (SPC-830, Becker and Hickl) for fluorescence lifetime measurements. For (3)’s green channel fluorescence intensity measurement mode, a fiber coupled 470nm LED (M470F3, Thorlabs) was collimated by L4, filtered by F3 (482/18, Semrock), and reflected by a removable mirror (M4); emission light was detected by a PMT (H7422-40, Hamamatsu). Alternatively, when fluorescence lifetime measurements were not needed, the bottom photometry unit was used. This simple “dual color fluorescence intensity photometry” unit consists of 470nm and 565nm LEDs (Thorlabs), two PMTs (H10770(P)A-40, Hamamatsu), and a Doric Minicube (FMC5_E1(465-480)_F1(500-540)_E2(555-570)_F2(580-680)_S, Doric Lenses) that are connected by patch cables. For both photometry units, LEDs were driven by a digital signal processor system (RX8-5-12, Tucker-Davis Technology) for frequency modulation to carry out locked in amplification of sensor signals detected by PMTs. In addition to the two main photometry units, a 593.5nm laser (SKU: YL-593-00100-CWM-SD-03-LED-F, Optoengine) and a 473nm laser (MBL-III-473, Optoengine) with independent patch cable connections were installed for Chrimson optogenetics and stGtACR2 optogenetics, respectively, for VTA DA neuron activity manipulation while monitoring NAc.

**Extended Data Fig. S2.**
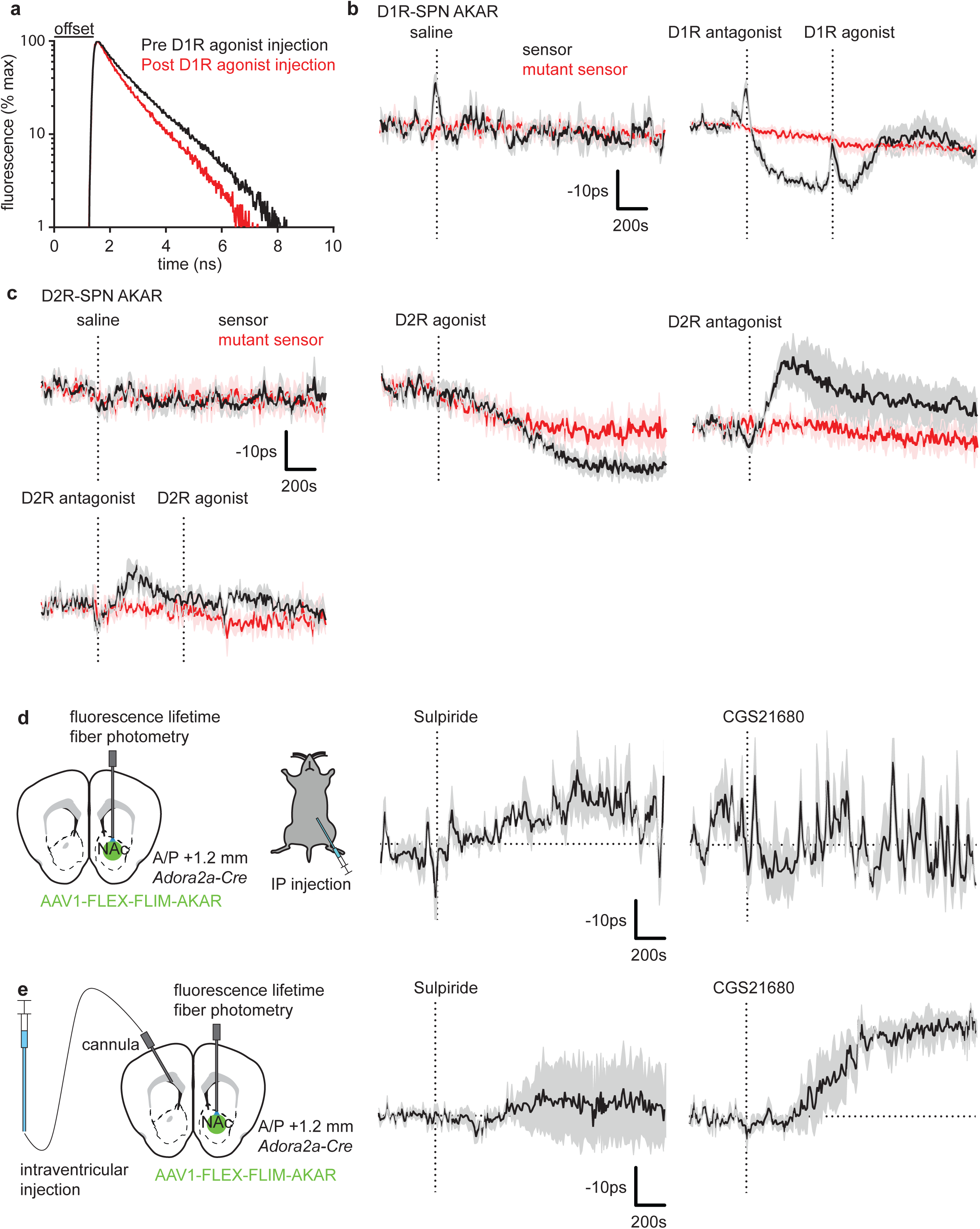
*In vivo* effects of drugs on SPN PKA activity. a) Fluorescence lifetime histogram of FLIM-AKAR expressed in D1R-SPNs of an example *Drd1a-Cre* mouse that was injected with D1R agonist. The black line shows the lifetime histogram measured before the injection during the baseline period (~60s before injection) whereas the red line shows the lifetime histogram measured after the injection at the time of the maximum lifetime change. The graph shifts leftward after D1R agonist injection demonstrating a faster decay in fluorescence due to the increased fraction of phosphorylated FLIM-AKAR sensors. b) D1R-SPN FLIM-AKAR responses of *Drd1a-Cre* mice to IP saline injection (n=4 mice) and IP D1R antagonist injection (n=7 mice) followed by IP D1R agonist injection (600s after D1R antagonist injection). c) D2R-SPN FLIM-AKAR responses of *Adora2a-Cre* mice to IP injection of various drugs. Top: saline response (*left*, n=5 mice for sensor and mutant sensor), D2R agonist response (*middle*, n=6 mice for sensor, 5 mice for mutant sensors), D2R antagonist response (*right*, n=6 mice for sensor, 5 mice for mutant sensor). Bottom: D2R antagonist followed by D2R agonist (n=5 mice for sensor and mutant sensor). d) D2R-SPN FLIM-AKAR responses of *Adora2a-Cre* mice to IP injection of drugs. *left*, Schematic of performing a fluorescence lifetime measurement using FLIM-AKAR while IP injecting a drug. *middle*, Sulpiride (D2R antagonist) IP injection (n=3 mice). *right*, CGS 21680 hydrochloride (A2AR agonist) IP injection (n=3 mice). e) D2R-SPN FLIM-AKAR responses of *Adora2a-Cre* mice to intraventricular (IV) injection of drugs. *left*, Schematic of fluorescence lifetime measurements of FLIM-AKAR while performing IV injection of a drug. Intraventricular cannula was implanted into a ventricle in the hemisphere opposite to the optical fiber implant. *middle*, Sulpiride (D2R antagonist) IV injection (n=3 mice). *right*, CGS 21680 hydrochloride (A2AR agonist) IV injection (n=3 mice). *For all graphs, dashed vertical line = end of injection, black=sensor (FLIM-AKAR), red=mutant sensor (FLIM-AKAR^T391A^). The solid line and shaded region show the average +/− SEM across mice

**Extended Data Fig. S3.**
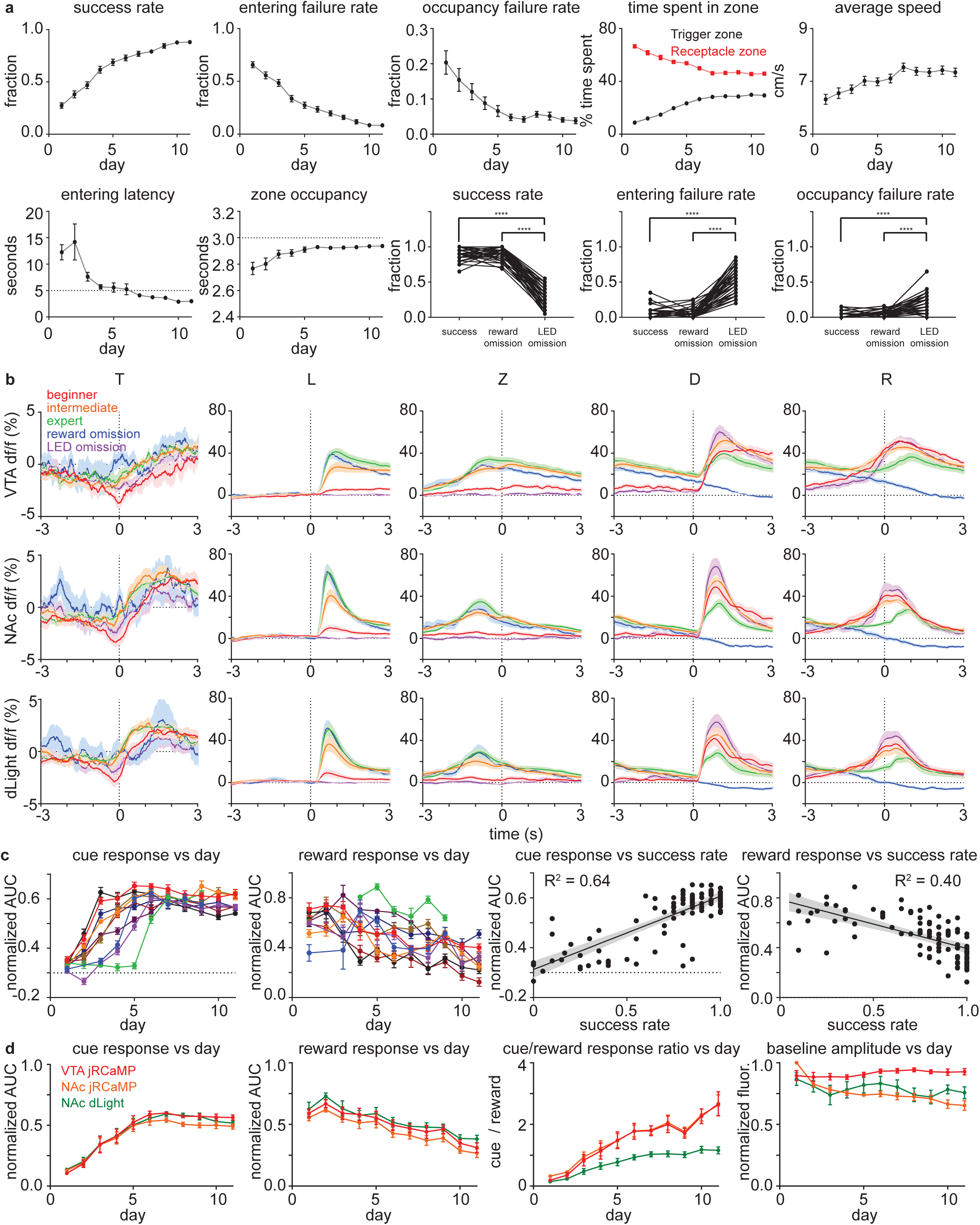
Plasticity in DA release and DA neuron activity dynamics is consistent with RPE. a) Behavioral parameters demonstrating that mice are able to learn the visual cue guided operant conditioning described in Fig. 2a. Top: from the left, success rate (number of rewarded trials / total number of trials), entering failure rate (number of receptacle zone entering failure trials / total number of trials), occupancy failure rate (number of premature receptacle zone exit trials / number of receptacle zone entering success trials), time spent in zone (time spent in a zone / total session time), average speed. Bottom: from the left, entering latency (delay to enter the receptacle zone after the LED cue), zone occupancy (time spent in the receptacle zone after entering the zone during a trial, 3s=maximum). Last three graphs depict success rate, entering failure rate, and occupancy failure rate comparisons for regular, reward omission, and rewarded LED omission sessions of expert mice. n=64 mice from all photometry behavior experiments. Error bar= SEM of averages across mice. b) DA neuron activity and DA release pattern across learning. The average responses for beginner, intermediate, expert, reward omission (of expert mice), and rewarded LED omission (of expert mice) trials are shown in red, orange, green, blue, and purple, respectively. Dashed vertical lines indicate the behavioral time stamps (T=trigger zone entry, L=LED on, Z=receptacle zone entry, D=pellet dispensing, R=receptacle entry). Top: df/f (%) of VTA jRCaMP signal showing VTA DA neuron soma activity. Middle: df/f (%) of NAc jRCaMP signal showing VTA DA neuron terminal activity. Bottom: df/f (%) of NAc dLight signal showing NAc DA level. Shaded area=SEM of averages across mice. n=10 mice. c) Patterns of cue and reward responses of dLight across training. From the left: LED cue responses of individual mice (different colors) measured by the area under the curve (AUC) of df/f (%) after cue onset ~ cue onset + 3s; reward responses measured by the AUC of 3s around the peak after receptacle entry; cue response (daily average) vs success rate for each mice; reward response (daily average) vs success rate for each mice. AUCs were normalized by the maximum value of each mouse across all sessions. Error bar=SEM of individual mouse average. Shaded area=95% confidence interval of the best linear fit. n=10 mice. d) Patterns of cue and reward responses across training for VTA jRCaMP, NAc jRCaMP, and NAc dLight are show in red, orange, and green, respectively. From the left: LED cue response as in panel c); reward response as in panel c); the ratio between the cue and the reward response; baseline (pre-trial) raw fluorescence estimating the change in a signal strength due to photo-bleaching and viral expression change. Both AUCs and raw fluorescence were normalized by the maximum value of each mouse across all sessions. Error bar=SEM of averages across mice. n=10 mice.

**Extended Data Fig. S4.**
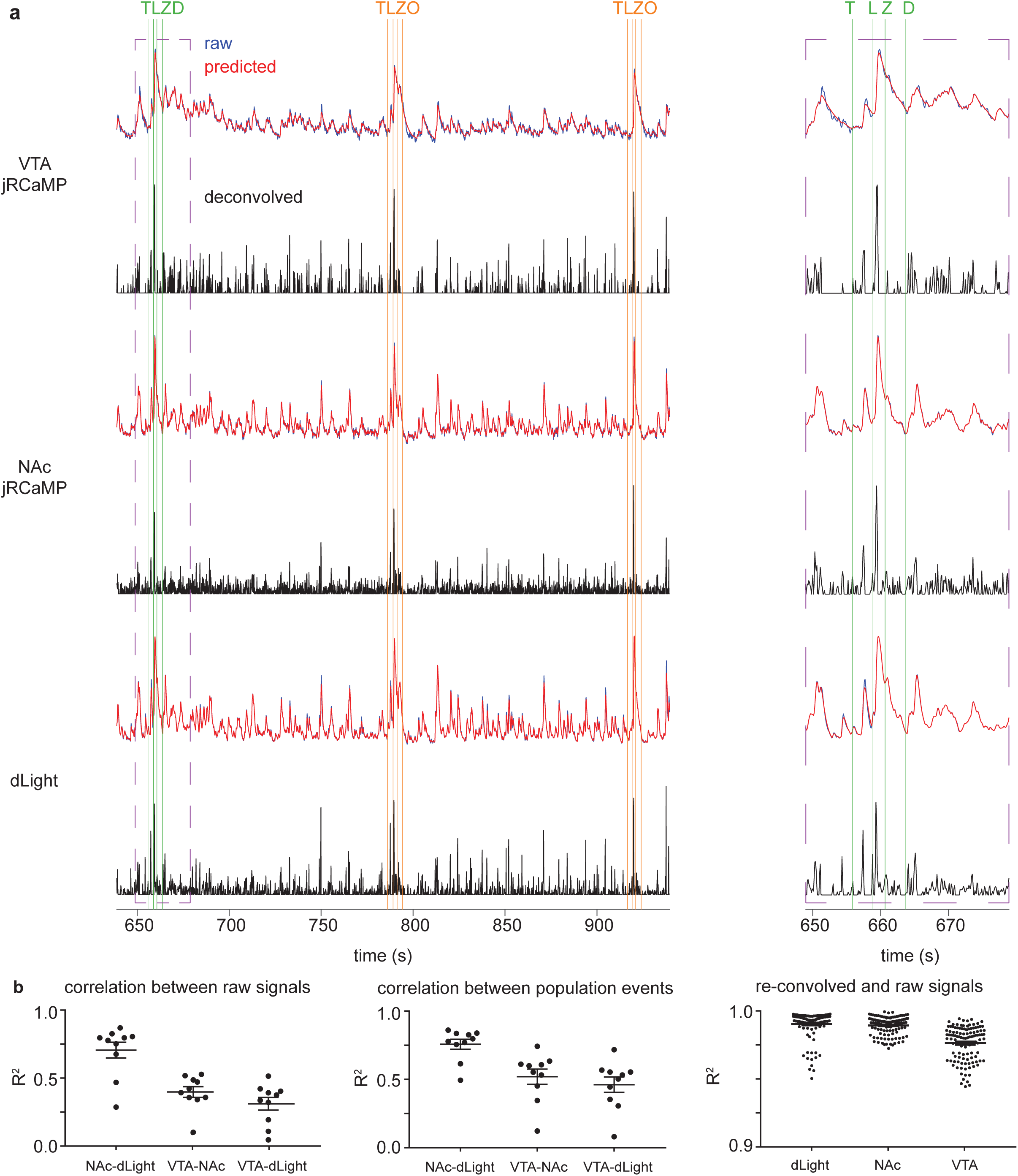
Deconvolution of photometry signals. a) VTA jRCaMP (top), NAc jRCaMP (middle), NAc dLight (bottom) signals from the portion of day 12 session of an example mouse, which includes 1 rewarded trial (green vertical lines) and 2 reward omission trials (orange vertical lines). For individual signal, actual fluorescence signals (blue) and fluorescence signals predicted from de-convolved populations events (red) were plotted on the top sub panel. Deconvolved population events (black) were plotted on the bottom sub panel. Purple dashed line box indicates the portion of the data that is shown on the inset on the right. Vertical lines indicate behavioral time points: T=trigger zone entry, L=LED on, Z=receptacle zone entry, D=pellet dispensing, and O=reward omission. b) Correlations between signals: NAc jRCaMP and NAc dLight signal correlation (NAc-dLight), VTA jRCaMP and NAc jRCaMP signal correlation (VTA-NAc), VTA jRCaMP and NAc dLight signal correlation (VTA-dLight). *left*, Correlation between signal amplitudes at all times. Error bar = SEM of averages across mice (n=10 mice). *middle*, Correlation between deconvolved fluorescence signals (population events). Error bar = SEM of averages across mice (n=10 mice). *right*, Correlation between the actual fluorescence signals and the fluorescence signals predicted from deconvolved populations events. Error bar = SEM across session values (n=115 sessions from 10 mice).

**Extended Data Fig. S5.**
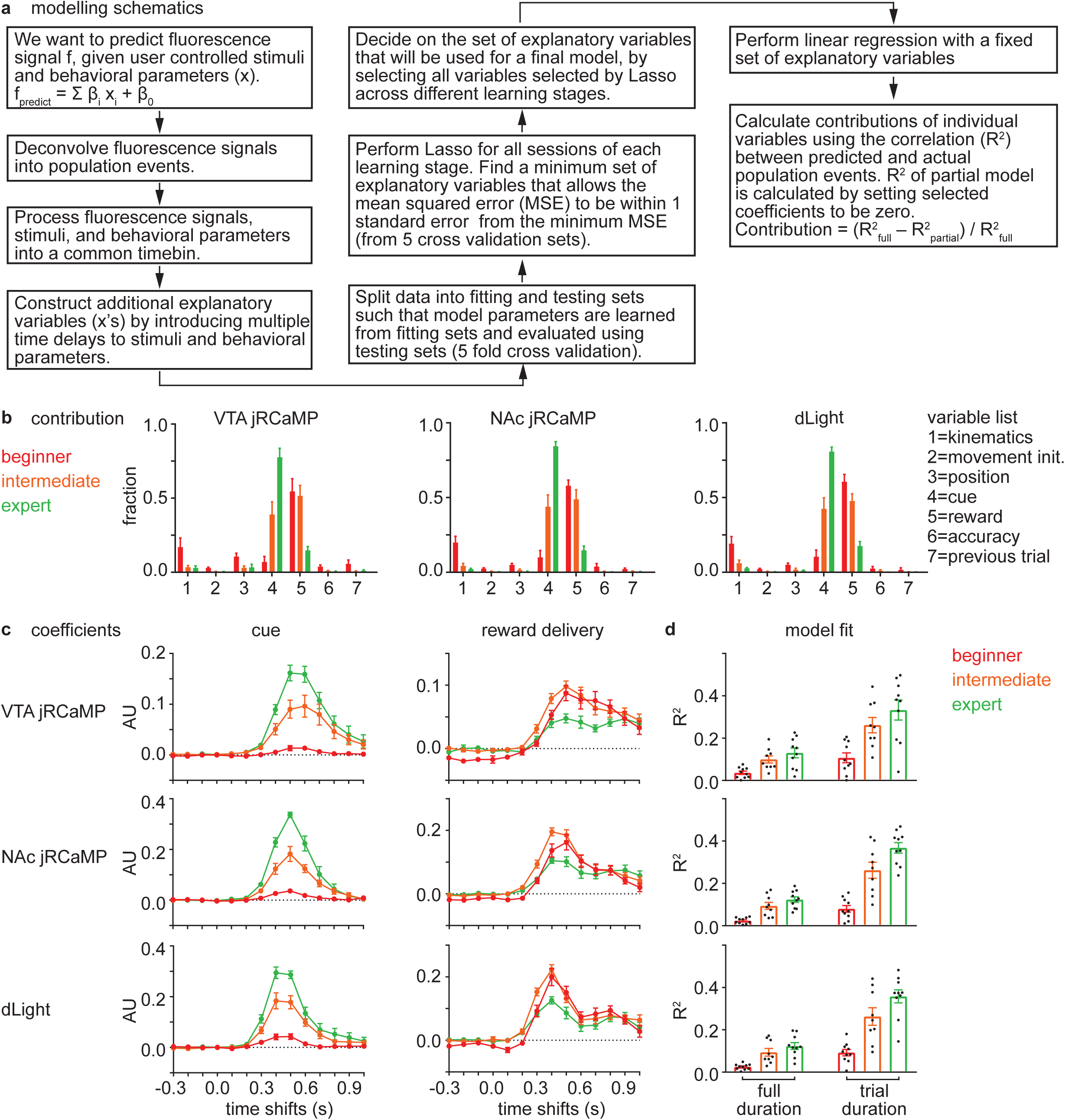
Relationship between DA neuron activity / DA release and behavioral parameters. a) Schematics of building a generalized linear model that relates user controlled stimuli and behavioral parameters to fluorescence signals. Briefly, there are 3 types of explanatory (independent) variables in the model. Continuous variables (speed, acceleration, rotation, position) continuously change their values as time passes. Event variables (movement initiation, cue, reward delivery, receptacle entry) are 0 except at a time point of an event when they temporarily change their value to 1. Whole trial variables (accuracy=0 for current trial failure, 1 for current trial success; previous trial=0 for previous trial failure, 1 for previous trial success) change their values in the beginning of a trial and stay constant until the next trial. b) Average variable contributions for VTA jRCaMP (*left*), NAc jRCaMP (*middle*), and dLight (*right*) for beginner, intermediate, and expert sessions shown in red, orange, and green, respectively. Contribution of each category was calculated by a method described in a). Kinematic variables include speed, acceleration, and rotation variables. Other categories are assigned to an individual variable (a set of time shifted variables). Error bar=SEM of averages across mice. n=10 mice. c) Average coefficient values for cue and reward delivery event variables (including all of the time shifted variables) for beginner, intermediate, and expert sessions shown in red, orange, and green, respectively. From the top, VTA jRCaMP model coefficients, NAc jRCaMP model coefficients, and dLight model coefficients. Error bar=SEM of averages across mice. n=10 mice. d) Average correlations between actual and predicted signals (population events) from the model for beginner, intermediate, and expert sessions shown in red, orange, and green, respectively. Left set of bars represent correlations during a full duration (−40~+80s respect to the trigger zone entry). Right set of bars represent correlations during a trial duration (−5+15s respect to the trigger zone entry). From the top, VTA jRCaMP model, NAc jRCaMP model, and dLight model. Error bar=SEM of averages across mice. n=10 mice.

**Extended Data Fig. S6.**
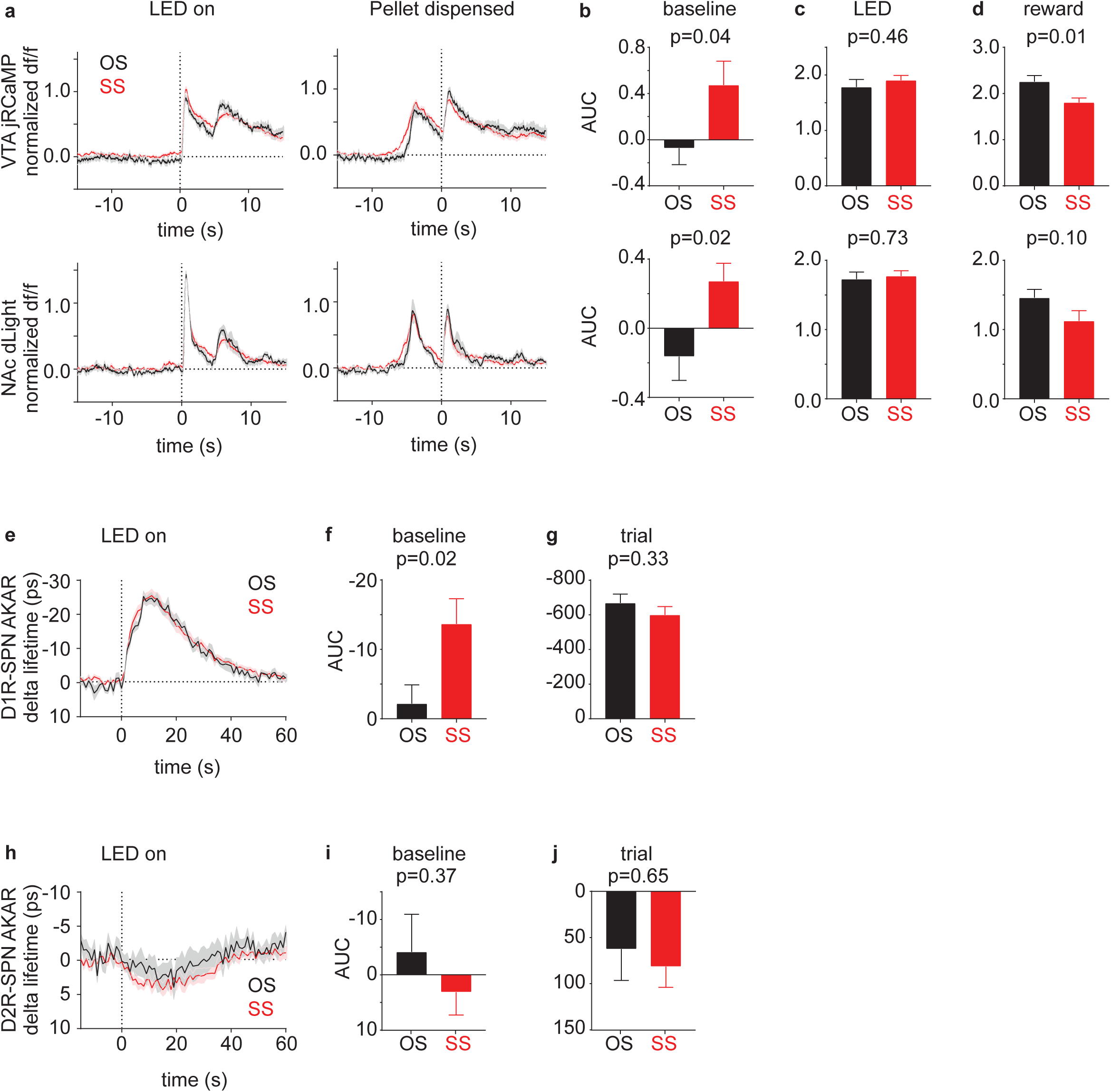
Trial history affects both baseline and peak amplitudes of DA neuron activity and DA release. To minimize the effect of photo-bleaching in analyzing the baseline period, we restricted our analysis to two consecutive trials. We separated consecutive trial pairs in the reward omission session (day 12) into those in which a successful reward-delivery trial followed a reward omission trial (OS=omission-success pair, black) or another reward delivery trial (SS=success-success pair, red). We found that DA increase in response to reward in the trial following a reward omission trial was significantly bigger than that in the trial following a rewarded trial (Extended Data Fig. 10a). In addition, the baseline DA level between the trials as measured before the LED cue was shifted higher following reward delivery than reward omission trial (Extended Data Fig. 10a), suggesting that expectation formed by trial history can have long-lasting effects (>=120s) on both baseline and peak DA level, consistent with previous findings^17^. Interestingly, the changes in baseline and peak size were also observed for DA neuron soma activity (Extended Data Fig. 10a). This suggests that DA neuron soma activity influences both phasic and tonic release of DA. Furthermore, the shift in baseline DA level was correlated with the change in baseline D1R-SPN PKA activity, suggesting that a change in baseline DA level does have a physiological effect on the downstream neuron. a) VTA jRCaMP (top) and dLight (bottom) response in *DAT-IRES-Cre* mice during the second trials of different trial pairs. *left*, normalized df/f (df/f normalized within the day 12 session for each mouse such that the 99 percentile response = 1) aligned to the time of LED cue. *right*, normalized df/f aligned to the time of pellet dispensing. df/f was calculated by using the average fluorescence of the reference period (− 20s~0s before the trigger zone entry) of the first trial as the f_0_. This allows the baseline period’s (−20s~0s before the LED cue) df/f of the second trial to be non-zero if the baseline period’s fluorescence of the second trial is different from that of the first trial. b) AUC of normalized df/f during the baseline period (−20s~0s before the LED cue). df/f was calculated as in panel a). c) AUC of normalized df/f between 0s~3s after LED cue. df/f was calculated by using the average fluorescence of reference period (−20s~0s before the trigger zone entry) of the second trial as the f_0_. d) AUC of normalized df/f between 0s~3s after pellet dispensing. df/f was calculated as in panel c). e) D1R-SPN FLIM-AKAR response in *Drd1a-Cre* mice during the second trials of different trial pairs. delta lifetime was calculated using the average lifetime of reference period (−20s~0s before the trigger zone entry) of the first trial as the reference value. f) AUC of delta lifetime of D1R-SPN FLIM-AKAR during the baseline period (−20s~0s before the LED cue). delta lifetime was calculated as in panel e). g) AUC of delta lifetime of D1R-SPN FLIM-AKAR during the trial (0s~30s after the LED cue). delta lifetime was calculated using the average lifetime of reference period (−20s~0s before the trigger zone entry) of the second trial as the reference value. h) D2R-SPN FLIM-AKAR response in *Adora2a-Cre* mice during the second trials of different trial pairs. delta lifetime was calculated as in panel e). i) AUC of delta lifetime of D2R-SPN FLIM-AKAR during the baseline period (−20s~0s before the LED cue). delta lifetime was calculated as in panel e). j) AUC of delta lifetime of D2R-SPN FLIM-AKAR during the trial (0s~30s after the LED cue). delta lifetime was calculated as in panel g). *Shaded area and error bar=SEM of averages across mice. n=10 mice (jRCaMP, dLight), 14 mice (D1R-SPN FLIM-AKAR), 18 mice (D2R-SPN FLIM-AKAR). For AUC comparisons, p-value was calculated from Welch’s t-test.

**Extended Data Fig. S7.**
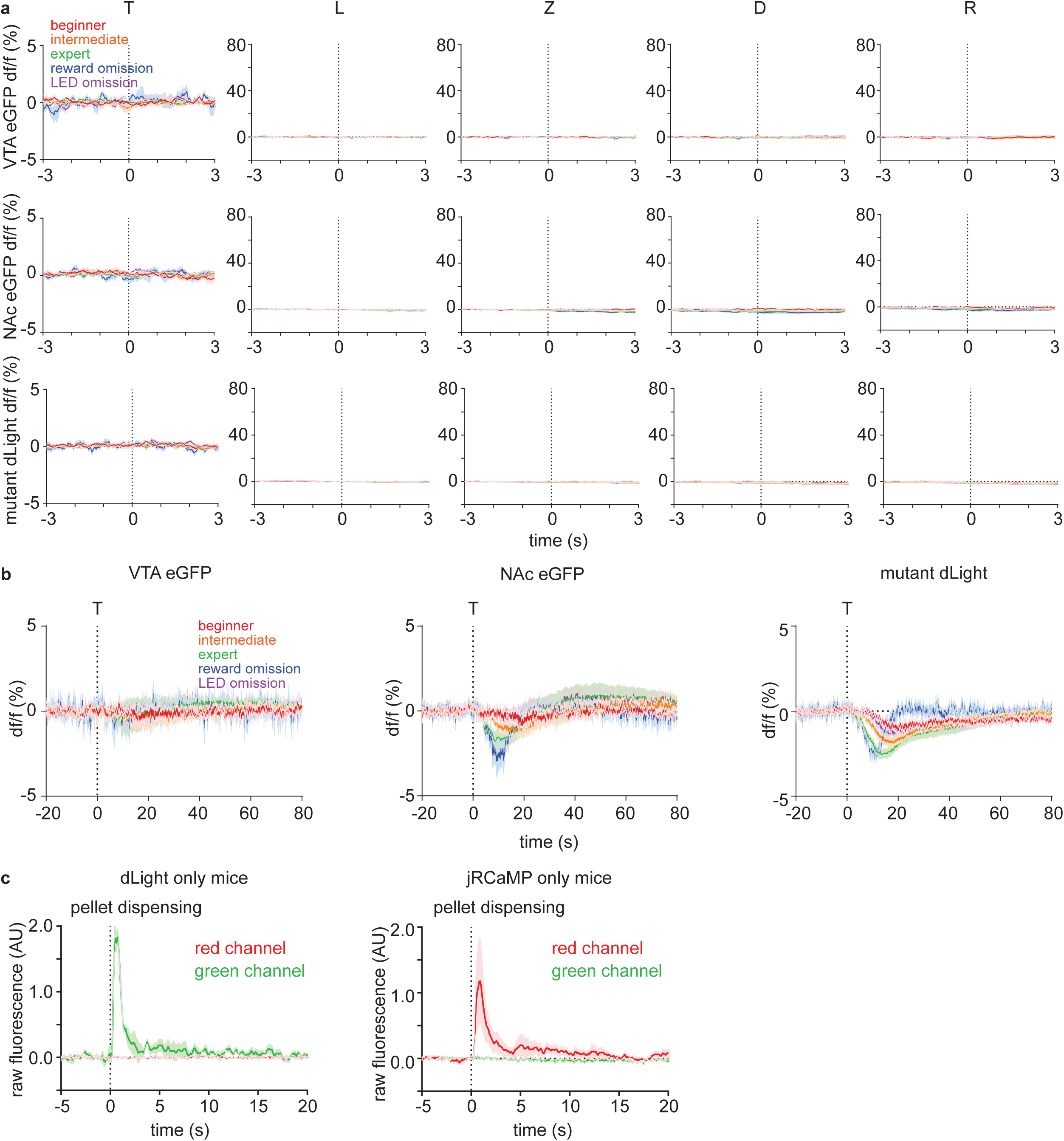
Movement and optical artifacts cannot explain dLight and jRCaMP signal patterns. a) df/f (%) of different controls. The average signals for beginner, intermediate, expert, reward omission (of expert mice), and rewarded LED omission (of expert mice) trials are shown in red, orange, green, blue, and purple, respectively. Dashed vertical lines indicate the behavioral time stamps (T=trigger zone entry, L=LED on, Z=receptacle zone entry, D=pellet dispensing, R=receptacle entry). Top: df/f (%) of eGFP signal from VTA of *DAT-IRES-Cre* mice (n=4 mice) that were injected with AAV1-Cag-FLEX-eGFP into VTA. Middle: df/f (%) of eGFP signal from NAc of *DAT-IRES-Cre* mice (n=4 mice) that were injected with AAV1-Cag-FLEX-eGFP into VTA. Bottom: df/f (%) of DA binding mutant dLight (D103A mutation) signal from NAc of *C57BL/6J* mice (n=8 mice) that were injected with AAV9-hSyn-dLight^D103A^ into NAc. b) df/f (%) of different controls that are magnified in df/f axis and de-magnified in time axis. VTA eGFP (*left*, n=4 mice), NAc eGFP (*middle*, n=4 mice), and NAc mutant dLight (*right*, n=8 mice) signal aligned to the time of trigger zone entry (dashed vertical line). There is a minor (comparable to sensor responses) but significant change in NAc eGFP and mutant dLight signal that develops across learning. c) Test for the optical crosstalk between green and red spectrum for simultaneous dual color photometry for dLight and jRCaMP. Mice were given unexpected free food pellets, and signal was aligned to the time of pellet dispensing. l*eft*, Raw fluorescence signal in red and green spectrum from NAc of *C57BL/6J* mice (n=3 mice, 10 trials/mouse) injected with AAV9-hSyn-dLight into NAc. *right*, Raw fluorescence signal in red and green spectrum from NAc of *DAT-IRES-Cre* mice (n=3 mice, 10 trials/mouse) injected with AAV1-hSyn-FLEX-NES-jRCaMP1b into VTA. *For all graphs, shaded area=SEM of averages across mice.

**Extended Data Fig. S8.**
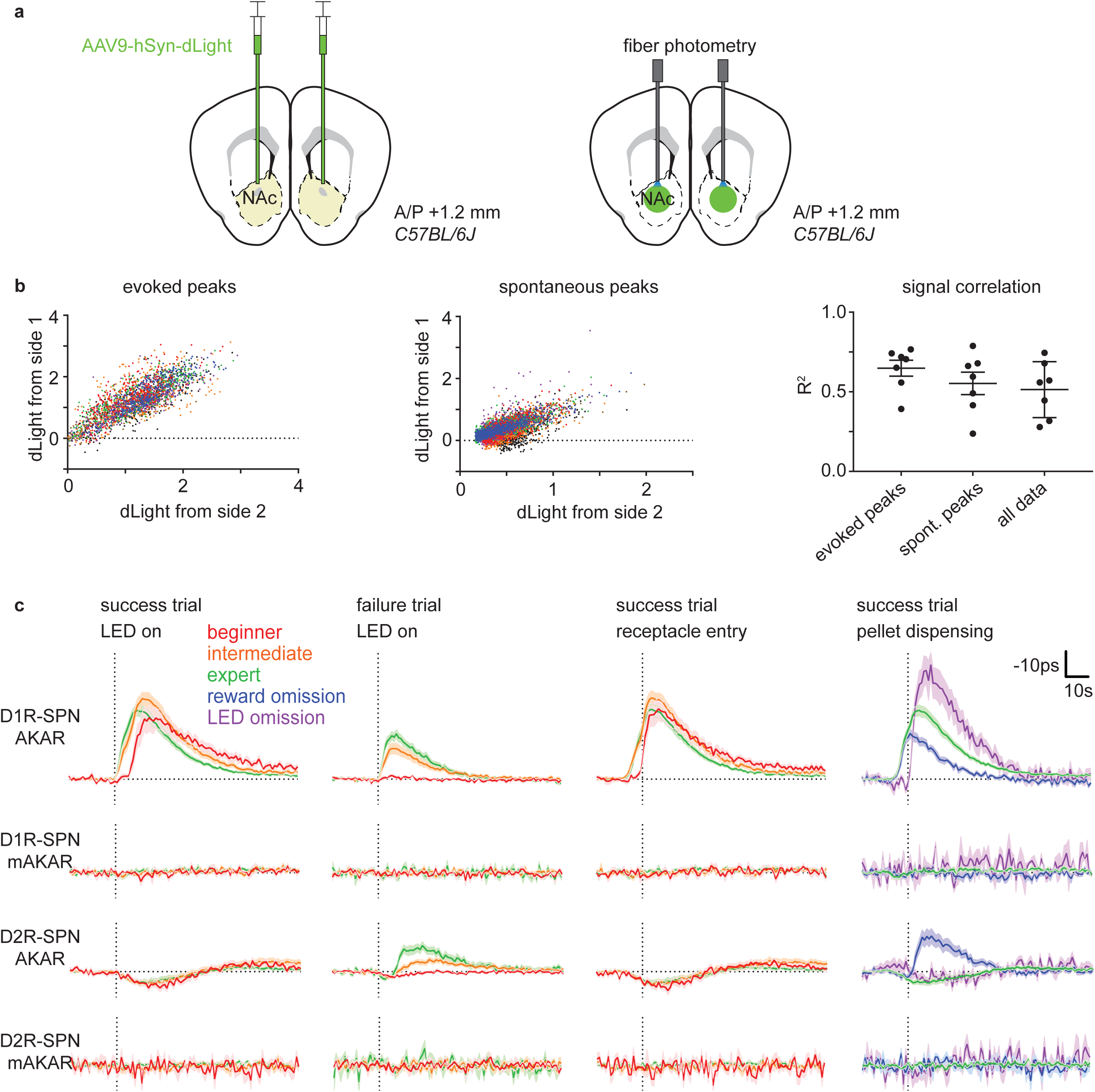
Bilateral dLight measurement and mutant FLIM-AKAR control experiments. a) Schematic describing a strategy to measure DA level in both hemispheres. AAV9-hSyn-dLight was bilaterally injected into NAc of *C57BL/6J* mice. Then, two optical fibers were implanted 200μm above the injection sites in two hemispheres. b) Relationship between dLight signals from two hemispheres. *left*, dLight signal (df/f normalized across all sessions for each mouse such that the 99 percentile response = 1) for LED cue and reward evoked peaks from two different sides for individual trials. Data points from the same mouse are plotted with the same color. *middle*, Normalized df/f of dLight signal for peaks during baseline period (−20s~0s before trigger zone entry) that rise above 2 STD of the baseline period. Plotted in the same way as the left. *right*, Correlation between bilateral dLight signals for evoked peaks, spontaneous peaks, and all data points. Error bar = SEM of averages across mice. n=7 mice. c) Comparison between FLIM-AKAR and FLIM-AKAR^T391A^, which has a point mutation at the PKA phosphorylation site, signals. AAV1-FLEX-FLIM-AKAR or AAV1-FLEX-FLIM-FLIM-AKAR^T391A^ was injected into NAc of *Drd1a-Cre* or *Adora2a-Cre* mice for these experiments. From the top, D1R-SPN FLIM-AKAR (D1R-SPN AKAR), D1R-SPN FLIM-AKAR^T391A^ (D1R-SPN mAKAR), D2R-SPN FLIM-AKAR (D2R-SPN AKAR), D2R-SPN FLIM-AKAR^T391A^ (D2R-SPN mAKAR). From the left, signals were aligned to the time (dashed vertical line) of “LED on” for success and failure trials separately, “receptacle entry” for success trials, and “pellet dispensing” for success trials. Signals for beginner, intermediate, expert, reward omission (of expert mice), rewarded LED omission (of expert mice) trials are shown in red, orange, green, blue, and purple, respectively. Shaded area=SEM of averages across mice. n=14 mice (D1R-SPN AKAR), 7 mice (D1R-SPN mAKAR), 18 mice (D2R-SPN AKAR), 6 mice (D2R-SPN mAKAR).

**Extended Data Fig. S9.**
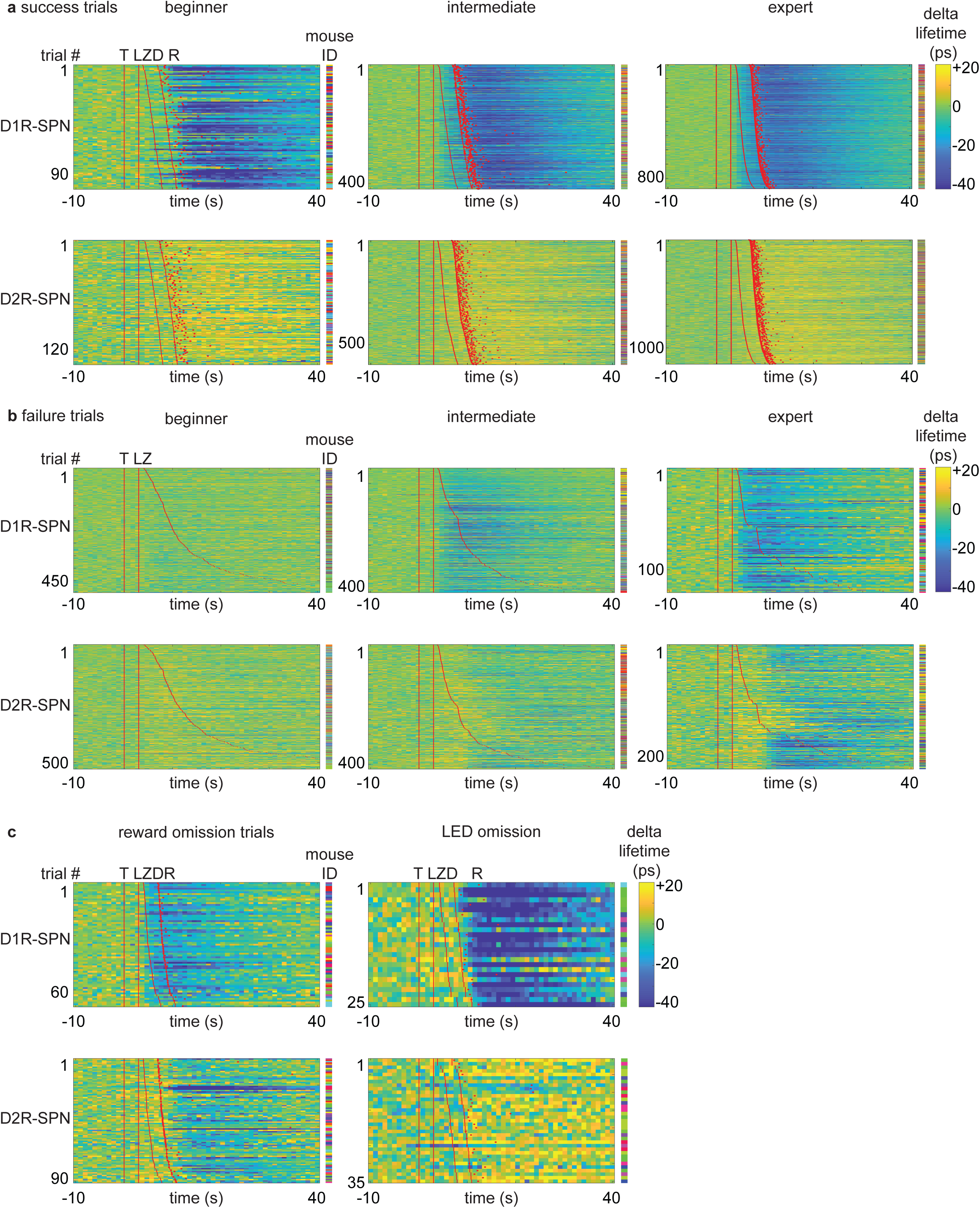
Plasticity in SPN PKA activity patterns during learning. a) Heatmaps of SPN FLIM-AKAR response for success trials during learning. Each row represents an individual trial. Red lines or dots indicate behavioral time stamps (T=trigger zone entry, L=LED on, Z=receptacle zone entry, D=pellet dispensing, R=receptacle entry). Different colors in mouse ID columns represent different mice for an individual row. Top: D1R-SPN FLIM-AKAR responses of *Drd1a-Cre* mice. n=98 trials (beginner), 418 trials (intermediate), 873 trials (expert) from 14 mice. Bottom: D2R-SPN FLIM-AKAR responses of *Adora2a-Cre* mice. n=134 trials (beginner), 596 trials (intermediate), 1152 trials (expert) from 18 mice. b) Heatmaps of SPN FLIM-AKAR response for failure trials during learning. Plotted as in panel a). Top: D1R-SPN FLIM-AKAR responses of *Drd1a-Cre* mice. n=497 trials (beginner), 402 trials (intermediate), 122 trials (expert) from 14 mice. Bottom: D2R-SPN FLIM-AKAR responses of *Adora2a-Cre* mice. n=528 trials (beginner), 416 trials (intermediate), 218 trials (expert) from 18 mice. c) Heatmaps of SPN FLIM-AKAR response for reward omission trials (*left*) and rewarded LED omission trials (*right*). Plotted as in panel a). Top: D1R-SPN FLIM-AKAR responses of *Drd1a-Cre* mice. n=69 trials (reward omission) from 14 mice, 25 trials (rewarded LED omission trials) from 6 mice. Bottom: D2R-SPN FLIM-AKAR responses of *Adora2a-Cre* mice. n=91 trials (reward omission) from 18 mice, 35 trials (rewarded LED omission trials) from 10 mice.

**Extended Data Fig. S10.**
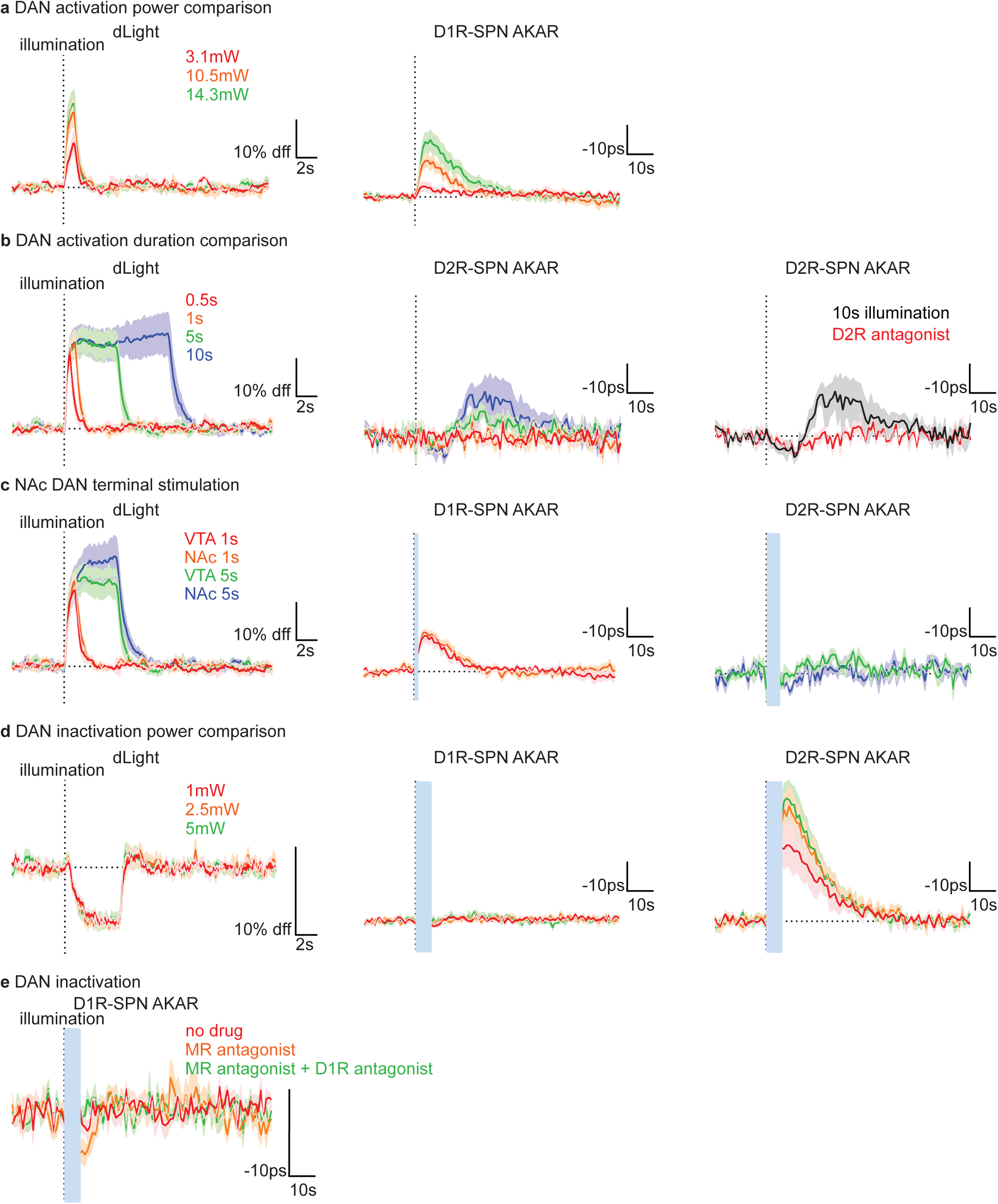
Transient change in DA neuron activity is sufficient to modulate SPN PKA activity in NAc. a) dLight (*left*, n=3 mice) responses in *DAT-IRES-Cre* mice and D1R-SPN FLIM-AKAR (*right*, n=6 mice) responses in *DAT-IRES-cre; Drd1a-Cre* mice to DA neuron activation (20Hz, 2ms pulse width, 1s illumination) for different levels of laser power (red=3.1mW, orange=10.5mW, green=14.3mW). b) *left*, dLight responses in *DAT-IRES-Cre* mice (n=3 mice) to DA neuron activation (20Hz, 2ms pulse width, 14.3mW illumination) for different durations of illumination (red=0.5s, orange=1s, green=5s, blue=10s). *middle*, D2R-SPN FLIM-AKAR responses in *DAT-IRES-Cre; Adora2a-Cre* mice (n=4 mice) plotted in the same way as the left. r*ight*, D2R-SPN FLIM-AKAR responses in *DAT-IRES-Cre; Adora2a-Cre* mice (n=4 mice) to 10s illumination without (black) and with (red) IP injection D2R antagonist at least 10mins before recording. c) dLight and SPN FLIM-AKAR responses to DA neuron terminal stimulation (20Hz, 2ms pulse width) in NAc (red=VTA DA neuron stimulation for 1s/10.5mW, orange=DA neuron terminal stimulation for 1s/7.7mW, green=VTA DA neuron stimulation for 5s/14.3mW, blue=DA neuron terminal stimulation for 5s/7.7mW). *left*, dLight responses in *DAT-IRES-Cre* mice (n=3 mice). *middle*, D1R-SPN FLIM-AKAR responses in *DAT-IRES-cre; Drd1a-Cre* mice (n=6 mice). *right*, D2R-SPN FLIM-AKAR responses in *DAT-IRES-Cre; Adora2a-Cre* mice (n=4 mice). d) dLight and SPN FLIM-AKAR responses to DA neuron inactivation (continuous illumination, 5s) for different levels of laser power (red=1mW, orange=2.5mW, green=5mW). *left*, dLight responses in *DAT-IRES-Cre* mice (n=4 mice). *middle*, D1R-SPN FLIM-AKAR responses in *DAT-IRES-cre; Drd1a-Cre* mice (n=7 mice). *right*, D2R-SPN FLIM-AKAR responses in *DAT-IRES-Cre; Adora2a-Cre* mice (n=5 mice). e) Revealing D1R-SPN FLIM-AKAR responses in *DAT-IRES-cre; Drd1a-Cre* mice to DA neuron inactivation (continuous illumination, 1mW, 5s) using muscarinic receptor antagonist. Average responses (n=4 mice) for no drug, IP injection of muscarinic receptor antagonist (Scopolamine hydrobromide) at least 10 mins before recording, IP injection of muscarinic receptor antagonist + D1R antagonist at least 10 mins before recording are shown in red, orange, and green. *For all graphs, shaded area=SEM of averages across mice, dashed vertical line=illumination onset. The average response of each mouse was calculated from 10 trials. Blue bars indicate the periods of blue laser illumination for stGtACR2 during which accurate FLIM-AKAR measurements were not possible.

**Extended Data Fig. S11.**
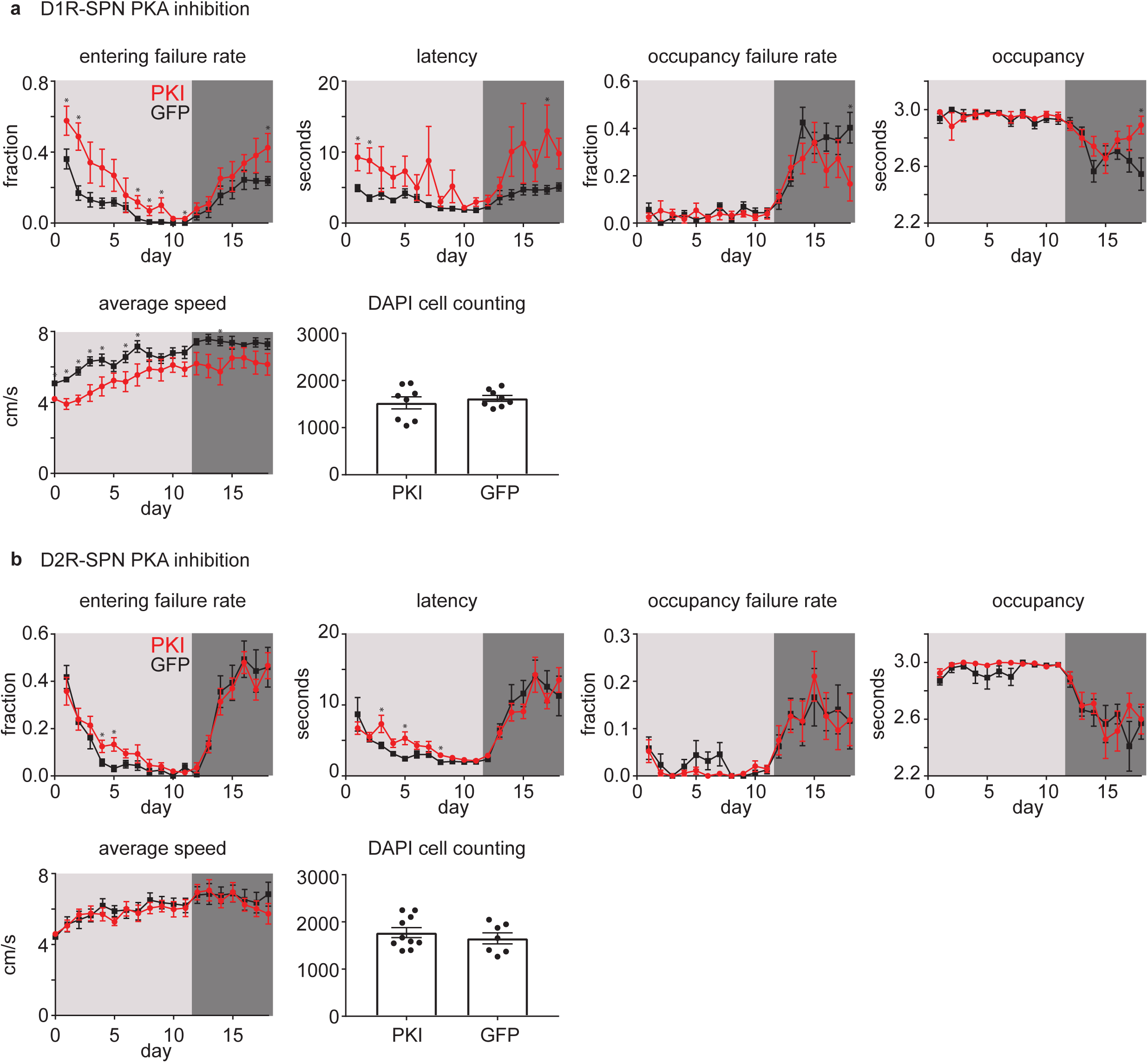
Selective PKA inhibition in SPNs partially impairs learning. a) Effect of D1R-SPN PKA inhibition. Top: from the left, entering failure rate (number of receptacle zone entering failure trials / total number of trials), latency (latency to enter the receptacle zone), occupancy failure rate (number of premature receptacle zone exit trials / number of receptacle zone entering success trials), and occupancy (time spent in the receptacle zone after entering the zone during a trial, 3s=maximum). Bottom: from the left, average speed and DAPI cell counting. PKI group (n=8 mice) and GFP group (n=8 mice) are shown in red and black, respectively. For all graphs, error bar=SEM of averages across mice, light gray=days of regular training, dark gray=days of extinction. * represents a p-value < 0.05 for Welch’s t-tests. b) As in panel a) for the effect of D2R-SPN PKA inhibition. n=10 mice (PKI) and 8 mice (GFP).

**Extended Data Fig. S12.**
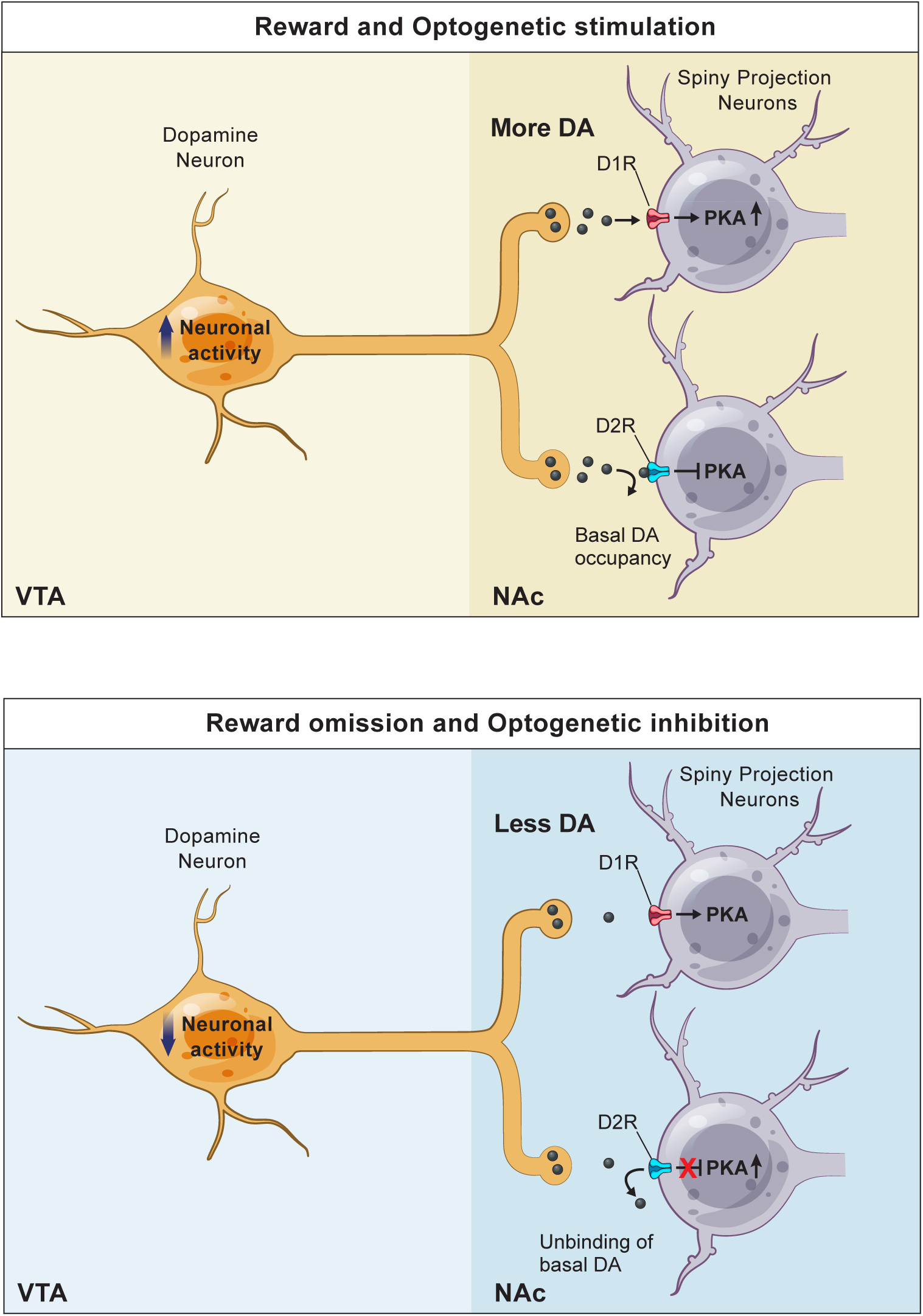
Model of DA action on SPN PKA activity. Overview of DA action on SPN PKA activity. **Top**: When DA neuron activity increases in response to a reward or optogenetic activation of DA neurons, more DA is released from the DA neuron terminals in the NAc. This increase in DA level allows DA to bind to D1R, which increases the activity of adenylyl cyclase, the level of cAMP, and ultimately the activity of PKA in D1R-SPNs. In contrast, the increase in DA level has a minimal impact on D2R, which is occupied by the basal level of DA. **Bottom**: When DA neuron activity decreases in response to a reward omission or optogenetic inhibition of DA neurons, DA release from the DA neuron terminals in the NAc decreases below the baseline. This decrease in DA level has a minimal impact on D1R, which is not occupied by the basal DA. In contrast, the decrease in DA level allows the basal DA to unbind from D2R, which disinhibits PKA activity in D2R-SPNs.

